# Temperature-dependent predation predicts a more reptilian future

**DOI:** 10.1101/2024.09.19.613816

**Authors:** John M. Grady, Jacob L. Amme, Kiran Bhaskaran-Nair, Varun Sinha, Samuel J. Brunwasser, Quentin D. Read, Sydne Record, Anthony I. Dell, Keith B. Hengen

**Affiliations:** Living Earth Collaborative, Washington University in St. Louis, St, Louis, MO, USA; Department of Biology, Washington University in Saint Louis, St. Louis, MO, USA; United States Department of Agriculture, Agricultural Research Service, Raleigh, NC, USA; Department of Wildlife, Fisheries, and Conservation Biology, University of Maine, Orono, ME, USA; National Great Rivers Research and Education Center, East Alton IL, USA

**Author notes:** These authors contributed equally to this work (John M. Grady, Jacob L. Amme). Last author (Anthony I. Dell, Keith B. Hengen).

## Abstract

Vertebrate diversity increases toward the tropics, but the extent to which this pattern varies with thermoregulatory strategy is unknown. A strong divergence, if confirmed, would imply a global restructuring of vertebrate communities with temperature. Here we present evidence for a novel latitudinal and thermal gradient of comparative vertebrate diversity. Synthesizing over 34,000 terrestrial species distributions, we observe a two-orders-of-magnitude shift in comparative richness with temperature, from endothermic dominance in temperate habitats and high elevations, toward parity with ectotherms in the lowland tropics. Next, we provide experimental support for an underlying thermal gradient of predation. Using machine vision tracking in over 4,500 endotherm-ectotherm predation trials, we show that thermally-mediated differences in performance favor endotherm predators in colder conditions and yield theoretically predicted outcomes, including strike count, distance traveled, and time to capture ectotherm prey. Finally, we integrate theory and data to forecast future patterns of diversity, revealing that as the world gets warmer, it will become increasingly reptilian.

---

One of the most widely recognized patterns in ecology is that species richness increases toward the tropics^1^, including vertebrate reptiles, amphibians, mammals, and birds^2^. What has received much less attention, however, is that the strength of this latitudinal pattern in diversity may deviate strongly with thermoregulatory status. In particular, ectothermic reptiles and amphibians — which rely on ambient heat for their body temperature — show a rapid increase in richness toward the tropics, whereas endothermic mammals and birds with stable body temperatures appear to have a more gradual rise^3,4^. Evidence is emerging that vertebrate abundance and predation shifts in a similar fashion. In the ocean, marine mammal abundance and total consumption rates are lowest in the tropics but increase rapidly toward the poles^5,6^. On land, a global synthesis found mammal abundance to be highest in temperate areas, while reptile abundance is greater in the tropics^7^. Similarly, Roslin et al. (2017) observed that the proportion of attacks on insects by mammals and birds rose in high latitudes and elevations, as attacks by arthropod predators declined^8^. Because large-scale and local-scale diversity are closely linked^9–11^, the potential implications are significant: a reorganization of vertebrate communities across the globe, from endotherm-dominance in high latitudes toward ectotherm-dominance in the tropics. However, much of the analyses on terrestrial richness and thermoregulation are taxon-specific and correlational, and a unifying model of comparative vertebrate diversity and its mechanisms is lacking.

What drives the shift from ectotherm to endotherm dominated communities? The decline in ectotherm diversity at high latitudes does not simply reflect a vulnerability to freezing — in the ocean, there is a strong, continuous shift with temperature toward endotherm dominance in the cold, despite waters only freezing near the poles^6^. Temperature-dependent predation has been proposed as a general mechanism for this global shift in marine vertebrate diversity toward endotherms in cold waters^5,6^. In particular, endothermic predators should be favored in cold environments where ectothermic prey metabolism is depressed and their movement becomes sluggish. This biotic advantage is formalized in Metabolic Asymmetry Theory (MAT), which advances predictions for predatory and competitive advantages on the basis of the asymmetric responses of endotherms and ectotherms to ambient temperature: ectotherms show an approximate exponential increase in performance with temperature until reaching a limit, but endotherm performance is invariant^6^. Ecosystem-scale support for MAT includes evidence that the fraction of prey consumed by marine endotherms increases in colder waters at a rate consistent with metabolic differences between endotherms and ectotherms^6^. Nonetheless, direct experimental support for MAT and temperature-dependent predation at the individual scale is still lacking. Does temperature shape the individual predatory success of endotherms under controlled conditions, and at rates consistent with theory? If confirmed, this would not only provide support for a general mechanism of comparative vertebrate diversity, but it would also offer insight into the consequences as well. More than half of all mammals and birds rely on ectothermic prey^12,13^. As global temperatures continue to rise, capturing cold-blooded prey will become more and more difficult, imperiling many endothermic species and shifting the balance of diversity toward ectotherms.

To address these questions, we first synthesize global datasets and quantify linkages between terrestrial vertebrate richness and thermoregulation. We demonstrate that the thermal sensitivity of comparative richness is virtually identical to the thermal sensitivity of metabolic rate, underscoring the role of metabolism in shaping global diversity. Next, we test core predictions of MAT and explore the mechanistic underpinnings of temperature-dependent predation using endothermic mice and ectothermic red runner cockroaches in controlled predation experiments. We find broad support for MAT, while also revealing how different behavioral components of predation are constrained by temperature. Finally, we integrate mechanism and macroecology to address a looming question: how will vertebrate diversity change as the planet warms?

## Temperature and Vertebrate Diversity

Both temperature and species diversity increase toward the tropics, but how they interrelate has been the source of considerable debate^14^. Using regional datasets, Allen et al. (2002) showed that ectothermic diversity increased at the same rate as metabolism for many taxa, suggesting deep linkages between physiology and species diversity^3^ (see also:^4,15–17^). Specifically, they showed that both metabolic rate *B* and richness *R* exhibit the same thermal dependence for many taxa:

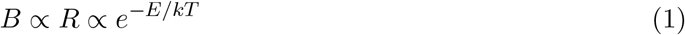

where ∝ indicates proportionality, *k* is Boltzmann’s constant, *T* is temperature in Kelvin and *E* is a thermal sensitivity coefficient (‘activation energy’) that is ~ 0.65 for metabolic rate^18^, corresponding to a ~ 2.5 fold increase per 10 °C (i.e., *Q*_10_ = 2.5).

Not all vertebrates are ectothermic, of course. On land, there are nearly as many endothermic mammals and birds as ectothermic reptiles and amphibians^2^. Consistent with their thermal stability, endotherm richness appear to have a weaker temperature dependence than ectotherm richness^3,4^. To provide a global assessment of thermal drivers of endotherm and ectotherm diversity, we synthesized the most extensive dataset of terrestrial vertebrate richness to date — spanning 34,366 species distributions of reptiles, amphibians, mammals, and birds. We quantified comparative patterns of diversity using the ratio of endotherm to ectotherm richness as a metric for relative composition (Fig. 1A). The pattern is clear: endotherms increase rapidly and systematically in comparative diversity toward the poles (Fig. 1B). This shift not only occurs with latitude but elevation as well (Fig. 1C), increasing threefold from 0 to 6000 m in tropical and subtropical habitats (− 30° to 30°, *p <* 0.001), as illustrated by the spike in relative endotherm diversity in the Himalayas and Rocky Mountains (Fig. 1A). An interesting secondary pattern is the decline in relative endotherm diversity in tropical deserts, between 15° and 30° north and south, driven by the elevated richness of squamates (snakes and lizards) in hot, unproductive habitats (Fig. 1A-B).

**Figure 1:**
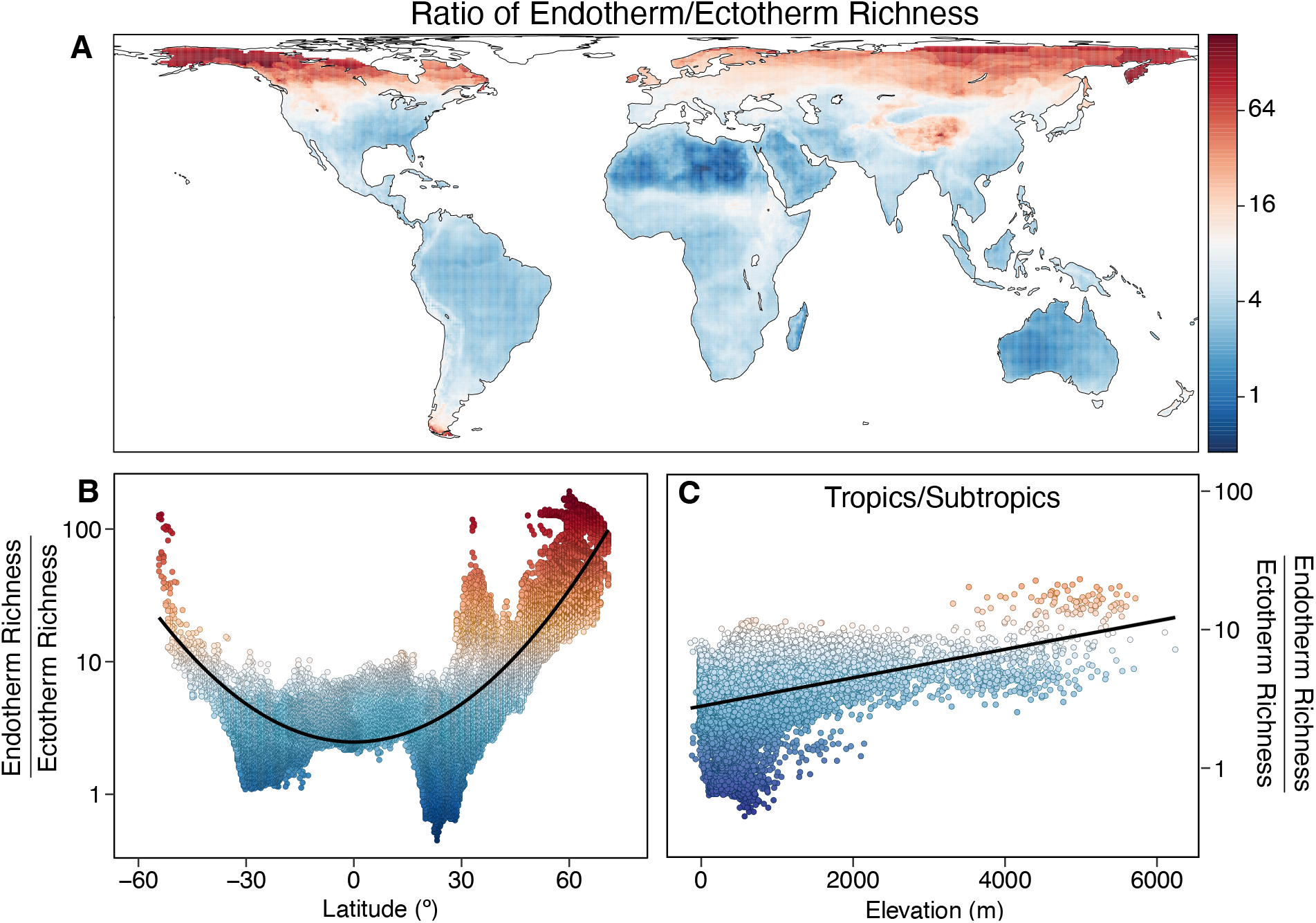
Comparative diversity of terrestrial endotherms and ectotherms. **(A)** Spatial map of the ratio of endotherm to ectotherm richness. Red indicates more endotherm species than ectotherms, blue indicates more ectotherm species than endotherms, and white is an equal number. **(B)** Relative endotherm richness increases toward the poles. **(C)** Relative endotherm richness increases with elevation. To control for thermal-latitudinal covariation, data was restricted to tropical and subtropical latitudes, from − 30° to 30°. Linear model: *y* = 1.0 + 0.00024*x, r*^2^ = 0.13, *p <* 0.001. Legend color in (A) applies to all panels. See also Figs. S1, S2.

The observed changes in vertebrate richness with latitude and elevation track changes in ambient temperature. Globally, richness in all major vertebrate taxa shows a positive but nonlinear association with temperature (Fig. 2A), but comparative patterns reveal a much more linear divergence (Fig. 2B). We observed an 85-fold shift in relative richness with temperature globally: in the coldest shared habitats, there are over 190 times more mammal and bird species compared to reptiles and amphibians, but in the warmest habitats, there are only 2.3 times as many. The magnitude of this compositional shift is almost identical to the thermal sensitivity of metabolic rate, with a ~ 2.5 fold increase in relative richness per 10 °C (Fig. 2B; spatial Bayesian hierarchical model: *E* = 0.64, *CI* : 0.63−0.64, *r*^2^ (fixed effects) = 0.79; linear model: *E* = 0.58, *CI* : 0.58−0.59, *r*^2^ = 0.80; see also Fig. S3). We also included mean annual precipitation, net primary productivity, and elevational range as predictors in spatial and linear models, but their relative contributions and effects on the slope were modest (linear model: temperature accounts for 89% of explained variance; log(precipitation) = 9%, log(NPP) = 1.2%, and elevation = 0.7%; Supp. Data 1, tabs 1−2).

**Figure 2:**
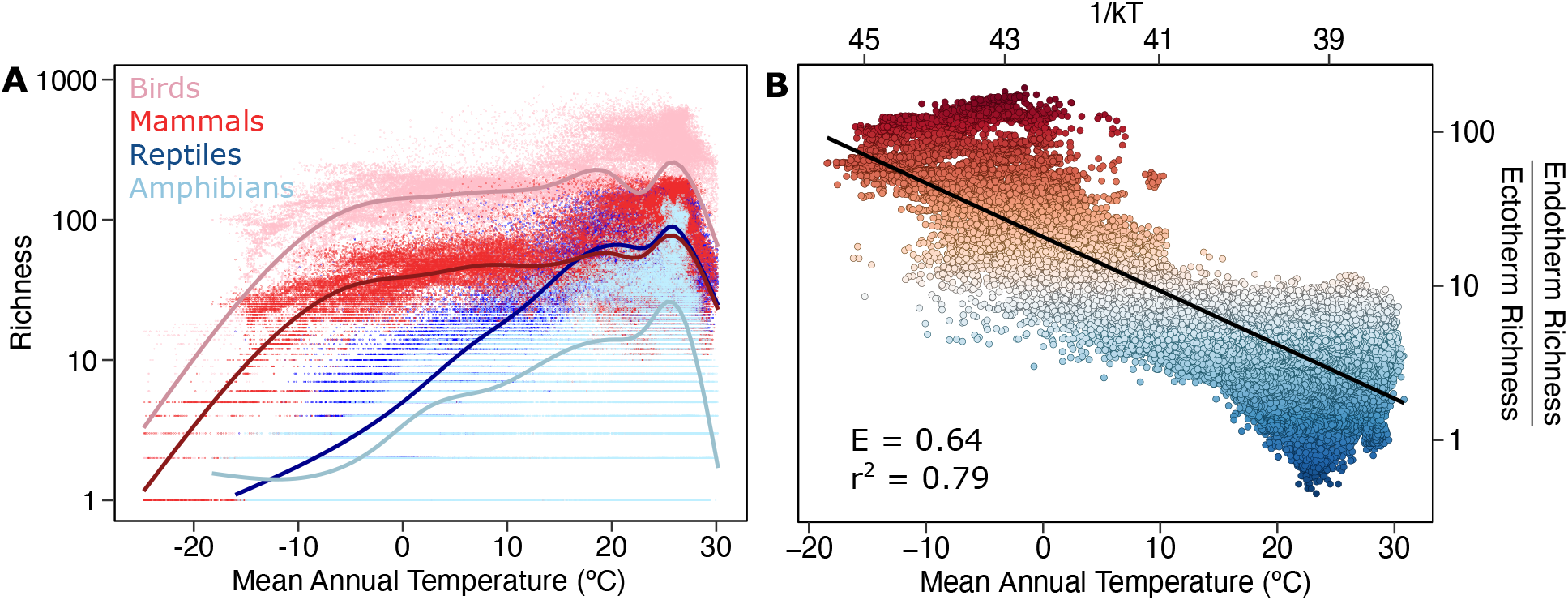
Temperature and global diversity of terrestrial endotherms and ectotherms. **(A)** All major taxa of vertebrates show a nonlinear increase in richness toward the tropics, but ectothermic amphibians and reptiles have a much steeper increase than endothermic mammals and birds. Endotherm diversity is relatively flat until ~ −5 °C mean annual temperature, or ~ 55° latitude, near the onset of boreal forests. **(B)** The ratio of endotherm to ectotherm richness values from (A) is plotted against surface temperature, revealing an approximately linear decline in comparative endotherm/ectotherm richness with temperature. The slope — where richness is regressed against inverse temperature (1*/kT*) — is indicated by the thermal sensitivity coefficient *E*. Note that the rate of decline in relative richness (Bayesian spatial model: *E* = 0.64) is almost identical in magnitude to the thermal sensitivity of metabolism (*E* = 0.65), suggesting that metabolic differences underlie global variation in comparative vertebrate diversity. See Fig. 1A for legend color.

This metabolic-based divergence in diversity extends beyond endothermy and ectothermy. We found additional evidence for the generality of this pattern within the largest terrestrial family of endothermic predators of ectotherms: shrews (Soricidae; *>* 400 spp)^19^. Shrews are highly diverse, invertivorous predators in the temperate northern hemisphere, but are largely absent from the lowland neotropics (Fig. S4A), ceding this invertebrate-rich habitat to slower metabolizing marsupials^20^. Only by undergoing daily torpor and depressing their basal metabolism and food requirements^21^ does an Old World subfamily of shrews claim this habitat in tropical Asia and Africa (Fig. S4A-B). Overlap between slow and fast metabolizing shrew subfamilies (Crocidurinae and Soricinae, respectively) does occurs throughout much of Eurasia and the ratio of their richness mirrors that of endotherms to ectotherms, with an identical thermal dependence to metabolism (Fig. S4C; *E* = 0.65, *CI* : 0.62 − 0.67, *r*^2^ = 0.61; see also Supp. Data 1, tabs 3−4). In this case of ecological competition, fast-metabolizing shrews act as the ‘endotherms’, whose energy-demanding lifestyle is supported by consuming cold and sluggish invertebrates. In the lowland tropics, however, where warm prey are too challenging to capture reliably, only ‘ectothermic’ shrews and marsupials are permitted.

## Testing Theoretical Mechanisms

A number of mechanisms have been proposed to explain the relationship between temperature and diversity, though few have focused on the role of thermoregulation. Metabolic Asymmetry Theory (MAT) proposes that temperature shapes predation and competition between endotherms and ectotherms via its asymmetric effect on metabolism, ultimately driving differential patterns of resource acquisition, abundance, and richness^6^. Faster metabolizing, endothermic species are better predators and competitors in the cold — where ectothermic prey are sluggish — but may struggle to acquire enough food where prey are warm and fast^22^. Specifically, a key assumption of MAT is that efficiency of prey escaping a motivated predator *ϵ* is proportional to their difference in speed *S*:

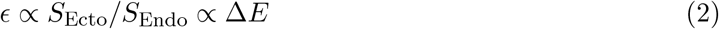

where Δ*E* ≡ *E*_SEcto_ − *E*_SEndo_. Thus, the escape efficiency for ectothermic prey from endothermic predator increases exponentially at warmer temperatures as its speed increases, shifting the favorability of species interactions toward ectotherms in warm habitats (see Methods).

Although ecosystem-scale evidence supports this model^5,6^, individual-scale evidence is limited, and its underlying mechanisms remain untested. Indeed, most previous thermal assessments of endotherm–ectotherm predation have focused on the consumption of slow-moving prey, where predatory rates are dictated by metabolic demand rather than the kinetic challenges of capturing warm, fast-moving targets (e.g., see references in^23^). More broadly, metabolic-based theories in ecology have been criticized for lacking mechanistic validation^24^. Unlike particles blindly colliding in space, predator and prey interactions involve nonrandom patterns of pursuit and escape that can dampen or exacerbate metabolic effects. Predators often exhibit learning and behavioral plasticity while foraging, further mitigating physiological constraints. Thus, it is unclear how closely predatory interactions track differences in metabolism with temperature.

To test MAT and evaluate the effects of temperature on endotherm–ectotherm predation, we manipulated ambient temperature and quantified pursuit and capture rates between an endothermic predator, the laboratory mouse (*Mus musculus*), and an ectothermic prey, the fast-moving, red runner cockroach (*Periplaneta lateralis*). In nature, many rodents are opportunistic predators of ectothermic invertebrates^25^, and research shows that even inbred laboratory mice with no prior hunting experience quickly learn to pursue and capture insects^26–29^. We trained C57BL/6 laboratory mice to hunt red runners at room temperature and then performed 4,747 individual foraging trials at five temperatures, ranging from 14 to 35 °C (Fig. S5). We assessed predator and prey movement using video tracking and machine vision-based markerless pose estimation, and evaluated the theoretical predictions of MAT at three interconnected levels: i) locomotion in isolation, ii) locomotion during pursuit/escape, and iii) foraging outcomes (Fig. 3, Fig. S5A-C). We also tested a secondary assumption of MAT — that predator learning can modulate thermal constraints on prey capture — and discuss how behavior overcomes, or fails to overcome, the kinetic constraints of temperature.

**Figure 3:**
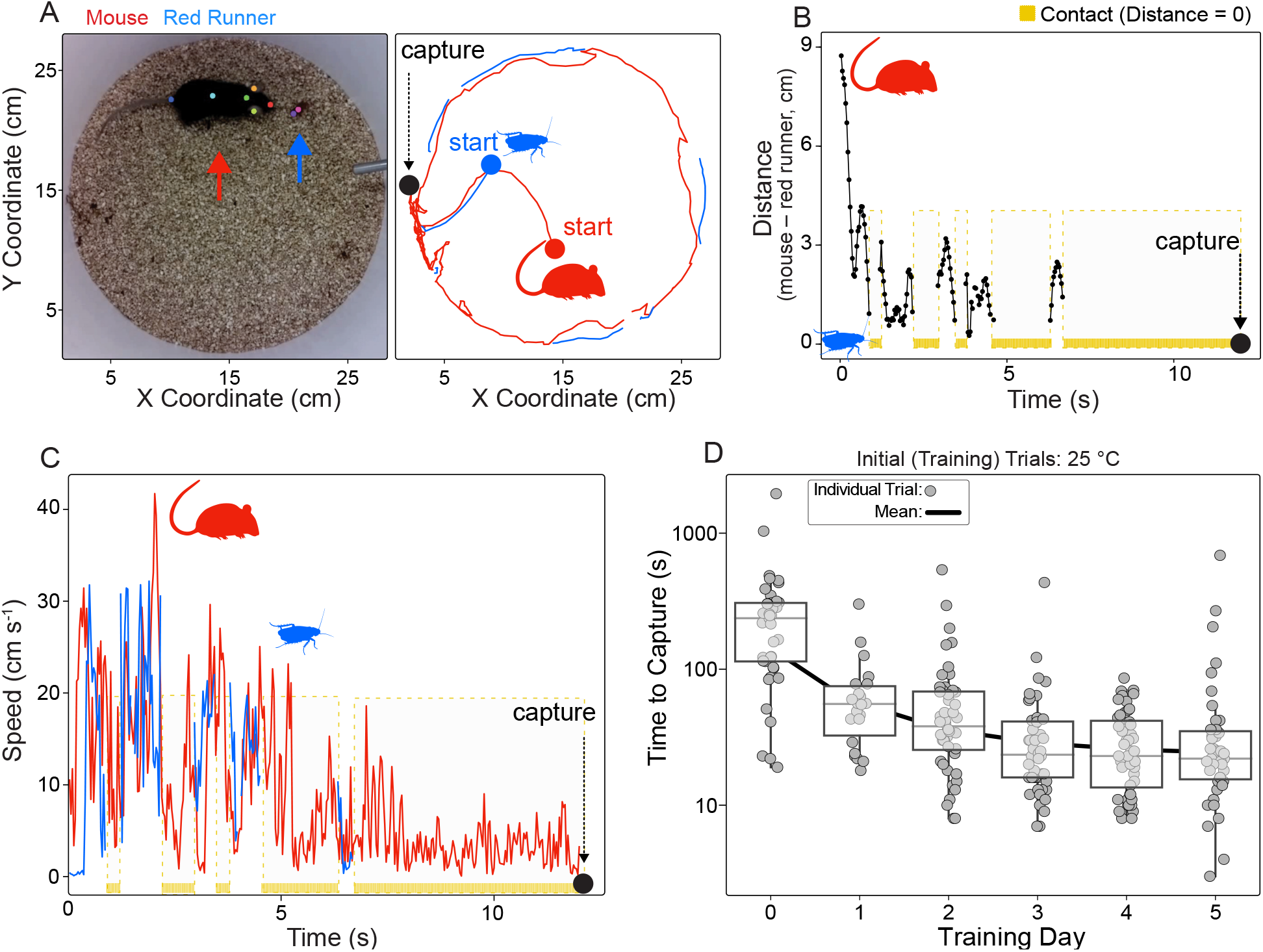
Laboratory mice rapidly learn to hunt and consume red runners. **(A)** (left) Video frame of mouse (red arrow) hunting a red runner cockroach (blue arrow). Colored points on the animals’ bodies illustrate the pose estimation of six points on the mouse and two points on the red runner. These coordinates were used to reconstruct animal paths (right), and to analyze components of foraging, including distance between predator and prey and contact frequency **(B)**, locomotory rates **(C)**, and the time to capture **(D)**. Mice rapidly approached asymptotic performance within three days of hunting trials (N = 8 animals).

## Experimental Tests of Theory

### Thermal Asymmetry of Locomotion

Mice rapidly learned to hunt during the initial, ten day trial stage, with the fitted time to capture declining from 331 s to 18 s (Fig. 3D, Fig. S6). After confirming that mice maintained a stable body temperature while red runners’ varied with thermal conditions (Fig. S7A-B), we measured their locomotion in isolation and tested model predictions (Fig. S5A). We expected that in isolation, mouse speed would be invariant to ambient temperature (*E* = 0), while red runners’ speed and acceleration would increase ~exponentially until reaching a peak, as described by *Eq. 2*. Across ectothermic species, the thermal sensitivity coefficient *E* has been found to be ~ 0.65 for respiration^18^, while in a synthesis of the thermal performance literature, Dell et al. (2011) found a slightly lower value for speed, with an average of *E* = 0.51 (*Q*_10_ = 2)^30^. Thus, we expected that the thermal sensitivity coefficient for red runner locomotion would be in the range of *E* = 0.5 − 0.65. To calculate *E*, we used mixed model linear regression, where temperature and red runner mass are fixed predictor variables, *E* is the slope, and mouse ID is a random effect (Fig. S8). Because red runner speeds typically peaked at 30 °C, we restricted regression fits to predation trials ranging from 14 – 30 °C (solid lines; Fig. 4), but we also plotted generalized additive model fits to show the full range across temperatures (dashed lines; Fig. 4).

**Figure 4:**
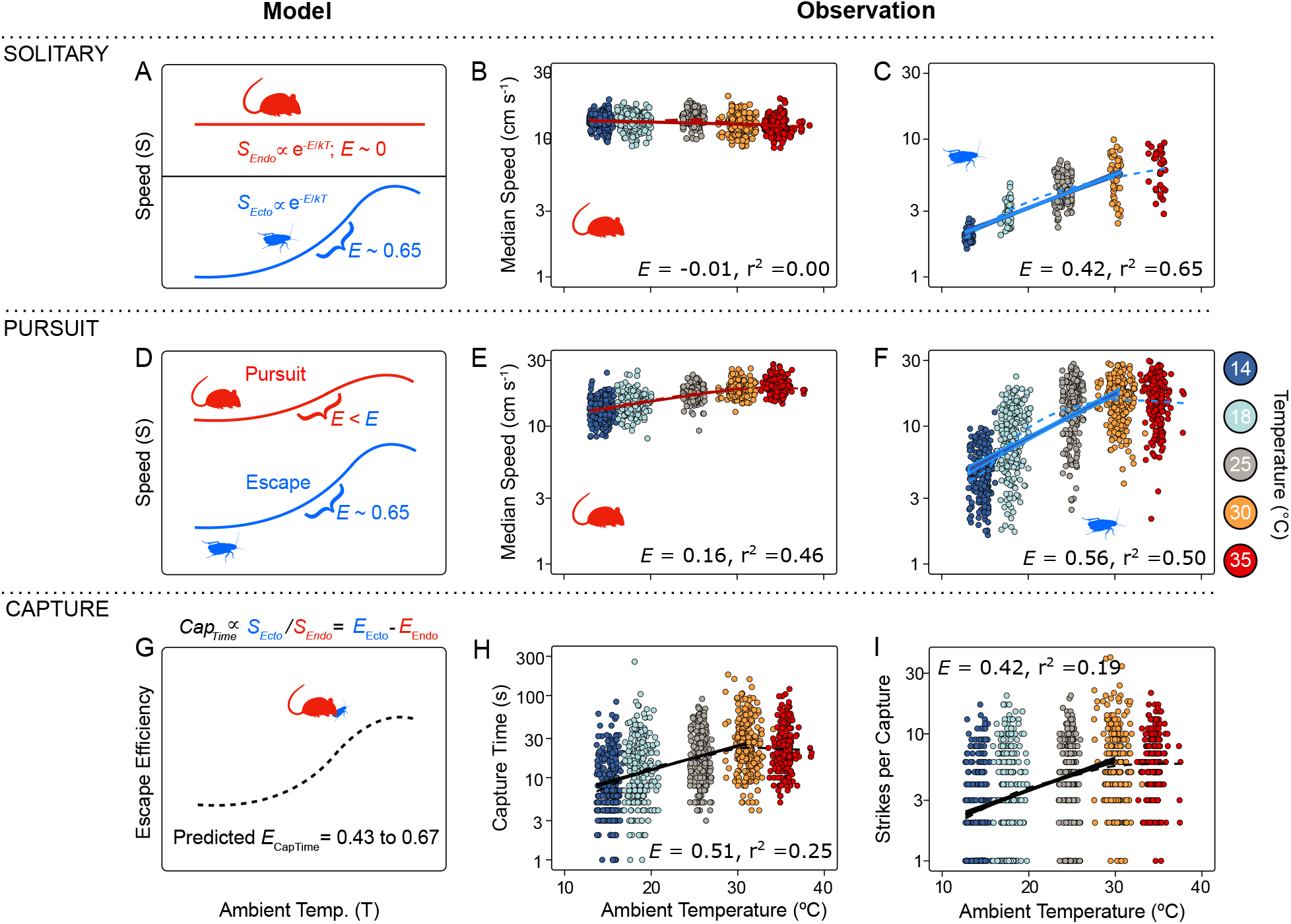
Foraging outcomes reflect thermal performance asymmetries. **(A)** In isolation, speed of endothermic mice should not depend on ambient temperature (*T*_*a*_), where the thermal sensitivity coefficient *E* ~ 0. In contrast, red runners are expected to increase speed at higher temperatures, where *E* ~ 0.5−0.65. **(B-C)** As predicted, mouse speed in isolation is invariant, while red runner speed increases with temperature (n mouse trials = 1107, n red runner trials = 227). **(D)** During pursuit, *E*_*SpeedMouse*_ is expected to be greater than 0 but less than *E*_*SpeedRRunner*_, as mice adjust their speed to track prey (0 *< E*_*SpeedMouse*_ *< E*_*SpeedRRunner*_); results support predictions **(E-F;** n mice = 1332; n red runner = 1191**). (G)** The time to capture is predicted to reflect the difference in thermal sensitivity of both maximum and median speeds (Δ*E*_*Speed*_) between mice and red runners in isolation (range: 0.43 − 0.67); **(H)** results support prediction (*E*_*CaptureTime*_ = 0.51, n = 1119). **(I)** Strikes per capture are similarly close (*E*_*Strikes*_ = 0.42, n = 1070), and the confidence interval includes the expected range. All *r*^2^ values represent main effects, see also Table 1, Data 1 Tab 5.

Consistent with predictions (Fig. 4A), mouse speed and acceleration in isolation showed little effect of temperature (Fig. 4B; *E*_*SpeedMouse*_ = − 0.0086, confidence interval (CI): − 0.017, − 0.00014; *E*_*AccMouse*_ = − 0.00048, CI: − 0.0072, 0.0063; see also Table 1, Supp. Data 1, Tab 5). As expected, red runner speed was thermally dependent and near the predicted range (Fig. 4C; *E*_*SpeedRRunner*_ = 0.42, CI: 0.38, 0.46), though the thermal dependence of acceleration was weaker (Supp. Data 1, Tab 5). Instantaneous, frame-to-frame locomotory rates followed the same pattern, with red runners showing higher ranges and central tendencies at warmer temperatures, while mice showed no change (Fig. S7C-D, Fig. S9). Note, all reported *E* values correspond to median rates, though estimates for maximum rates are similar (Table 1).

**Table 1:** Thermal sensitivities (*E*) of locomotion and hunting behavior. *E* represents the thermal sensitivity coefficient. Since distance traveled is the product of speed and time, *E*_*D*_*istance* is expected to be higher than either component. *r*^2^ Main corresponds to *r*^2^ associated with the main predictor variables, temperature (1/kT) and red runner mass. *r*^2^ Full is for the main effects plus mouse ID as a random effect. — indicates not applicable.

| Thermal Rates | $E_{Predicted}$ | $E_{Observed}$ | CI | $r^2$ Main | $r^2$ Full |
| --- | --- | --- | --- | --- | --- |
| ***In Isolation*** |  |  |  |  |  |
| *Speed (Median)* |  |  |  |  |  |
| Red Runner | 0.5-0.65 | 0.42 | 0.38, 0.46 | 0.65 | — |
| Mouse | 0 | -0.0086 | -0.017, -0.00014 | 0.0031 | 0.14 |
| *Speed (Maximum)* |  |  |  |  |  |
| Red Runner | 0.5-0.65 | 0.64 | 0.59, 0.70 | 0.75 | — |
| Mouse | 0 | -0.030 | -0.048, -0.011 | 0.0082 | 0.098 |
| ***In Pursuit*** |  |  |  |  |  |
| *Speed (Median)* |  |  |  |  |  |
| Red Runner | 0.5-0.65 | 0.56 | 0.53, 0.60 | 0.50 | 0.52 |
| Mouse | < 0.56 | 0.16 | 0.15, 0.17 | 0.46 | 0.47 |
| *Speed (Maximum)* |  |  |  |  |  |
| Red Runner | 0.5-0.65 | 0.62 | 0.60, 0.65 | 0.65 | 0.68 |
| Mouse | < 0.62 | 0.27 | 0.25, 0.29 | 0.37 | 0.45 |
| Mouse-Red Runner Distance | — | 0.0036 | -0.049, 0.056 | 0.015 | 0.11 |
| ***Strike*** |  |  |  |  |  |
| Strikes per Capture | — | 0.42 | 0.38, 0.47 | 0.19 | 0.46 |
| Strike Rate | — | -0.040 | -0.0075, -0.0048 | 0.011 | 0.044 |
| ***Capture*** |  |  |  |  |  |
| Time per Capture | 0.43-0.67 | 0.51 | 0.47, 0.55 | 0.25 | 0.56 |
| *Dist. per Capture* |  |  |  |  |  |
| Red Runner | > 0.51 | 1.4 | 1.3, 1.5 | 0.47 | 0.62 |
| Mouse | > 0.51 | 1.1 | 1.0, 1.1 | 0.46 | 0.62 |

### Predation Across Temperatures

Next, we conducted hunting trials at different temperatures following the same experimental protocol. Similar to their rates in isolation, we predicted that during pursuit red runners would exhibit an approximately exponential increase in locomotion with temperature until reaching a peak. We also expected that pursuing mice would show a positive but flatter response to temperature as they tracked their prey (i.e., 0 *< E*_*Mouse*_ *< E*_*RedRunner*_; Fig. 4D). Results supported predictions: red runner speed increased rapidly with temperature until reaching a plateau at 30 °C (*E*_*SpeedRRunner*_ = 0.56; CI: 0.53, 0.60), while mice showed a positive and weaker thermal sensitivity during pursuit (Fig. 4E-F; *E*_*SpeedMouse*_ = 0.16, CI: 0.15 − 0.17; see also Fig. S7E-F).

### Testing Metabolic Asymmetry Theory

According to MAT, predatory outcomes are theorized to reflect the difference in thermal sensitivity between predator and prey, where *E*_Ecto_ − *E*_Endo_ ≡ Δ*E*. Specifically, a key prediction of MAT is that the difference in thermal sensitivity of locomotion between a predator and its prey (Δ*E*_Speed_) determines the efficiency of escape *ϵ* (i.e, escape per encounter; *Eq. 2*). Since red runners could not exit the hunting arena, we used time to capture as a proxy for escape efficiency, as longer pursuit times increase the likelihood that prey find refuge or outpace their pursuer in nature. To predict the thermal sensitivity of time to capture (*E*_*CaptureTime*_), we used two measures of Δ*E*_*Speed*_ observed in isolation — median and maximum speed — as both variables are likely relevant to predation^31^ (Fig. 4B-C; Table 1). Δ*E*_*Speed*_ ranged from 0.43 − 0.67, so we predicted *E*_*CaptureTime*_ to fall within this range (Fig. 4G). Results uphold predictions: *E*_*CaptureTime*_ = 0.51 (Fig. 4H; CI: 0.47 − 0.55, *r*^2^ = 0.56; see also Table 1). Thus, we find support for the core MAT prediction that differences in thermal sensitivity drive the the ability of prey to escape predators. In addition, we found that *E*_*CaptureTime*_ was comparable in magnitude to the thermal sensitivity of strikes required to subdue prey: *E*_*Strikes*_ = 0.42 (Fig. 4I; CI: 0.38 − 0.47, *r*^2^ = 0.46; see also Table 1). This indicates that the challenge of capturing prey at higher temperatures is not merely in catching up to them, but in subduing metabolically vigorous animals. These thermal hunting outcomes occurred consistently, both in pooled results (Fig. 4), and for individual mice (Fig. S10).

### Behavior and Thermal Constraints

Learning and behavioral plasticity play key roles in predation and have the potential to buffer against environmental effects, including temperature-driven shifts in prey behavior^32^. Nonetheless, in our experiments, hunting outcomes remained tightly linked to thermal locomotion asymmetries observed in isolation, closely matching MAT predictions (Fig. 4). To understand how learning modulates but fails to overcome thermal constraints, we examined predation performance at three phases: pursuit, striking, and capture. In all three phases, performance improved rapidly with learning during initial trials at room temperature (Fig. 5, left column), after which thermal trials began (Fig. 5, right column).

**Figure 5:**
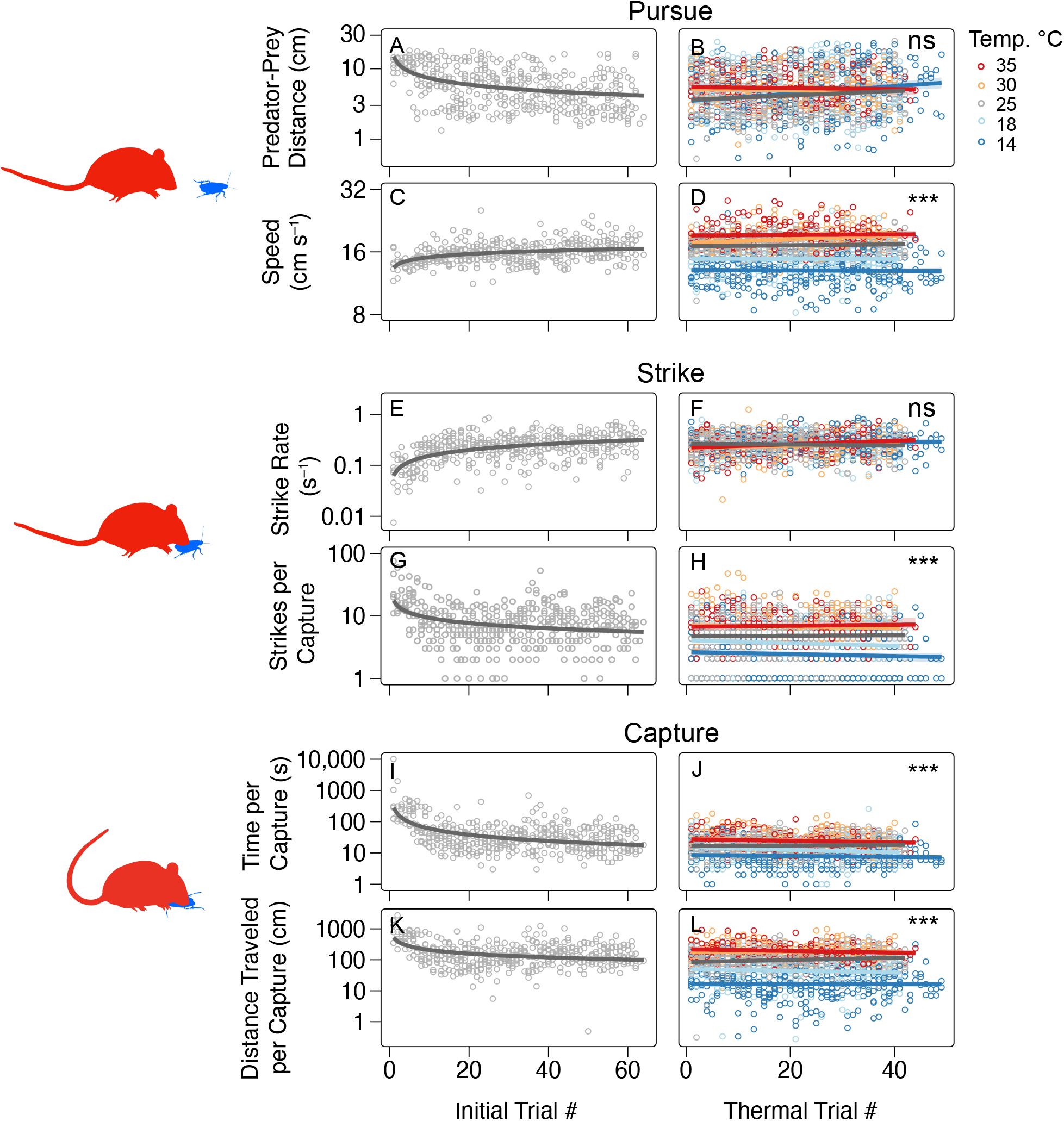
Learning and thermal constraints on predatory behavior. **Left column:** Mice showed improved performance over the course of initial trials, conducted at room temperature (25 °C). **(A, C)** As mice learned to pursue prey, they increased their speed and decreased their distance from prey. Strike rate increased with the number of trials **(E**) while strikes per capture declined over time as mice became more efficient at hunting **(G)**. Over the course of initial trials, the time **(I)** and distance traveled **(K)** needed to capture prey declined. For regression fits, log response was regressed against log trial number. **Right column**: thermal trials followed the initial trial period. **(B, D)** The average distance from prey remained constant across temperature regimes, as mice adjusted their speed to keep up with faster-moving prey in warmer conditions. **(F, H)** Strike rate also remained consistent across temperatures, although under warmer conditions, more strikes were required to subdue prey. **(J, L)** Ultimately, the time and distance traveled to capture prey showed a strong thermal signal. For regression fits, log response was regressed against trial number. See Data 1, Tabs 6-7 for results.

During the pursuit phase, mice quickly increased their speed and closed in on prey as they learned to track red runners (Fig. 5A,C). Since red runner speed was temperature-dependent, pursuing mice adjusted accordingly, maintaining a thermally-invariant distance of 4.6 cm throughout the trials (Fig. 5B,D; *p* = 0.08). Likewise, in the strike phase, mice struck at consistent rates across temperatures (Fig. 5F; *p* = 0.07), averaging one strike every 3.9 s. Thus, mice compensated for shifts in prey behavior with temperature, resulting in thermal invariance of both prey distance and strike rate.

Beyond these adjustments in locomotion, mice may have also learned to anticipate prey behavior across thermal conditions. In normal thermal trials, the temperatures of both the prey and hunting arena were matched, but in subsequent trials we mismatched them by placing warm prey in cold arenas and cold prey in warm arenas. These thermal mismatches consistently altered hunting outcomes even though predator and prey locomotion remained unchanged, suggesting that mice use thermal cues to anticipate prey before their introduction (Fig. S11; Cohen’s D: 0.5 − 1.1; see also Supp. Data 1, Tab 8).

Crucially, behavioral adjustments by mice had a limited impact compared to the dominant effect of temperature on hunting performance. Overall, prey were much harder to subdue at higher temperatures: the number of strikes required to incapacitate prey more than doubled, from 2.4 at 14 °C to 6.1 at 35 °C (Fig.5H; p *<* 0.001). As a result, the time and distance required to capture prey increased 3 to 11-fold respectively per 11 °C (Fig.5J,L). Ultimately, faster speed and reduced vulnerability of prey to capture at higher temperatures overwhelmed any compensatory strategies employed by mice, generating the strong thermal signal in predation predicted by theory.

### Forecasting Global Vertebrate Diversity

Using theory and data, we have shown that endothermic predators face increasing difficulty in capturing ectothermic prey at warmer temperatures. This not only poses challenges for endothermic predators but also provides significant advantages to ectothermic prey and competitors. Thus, we expect rising global temperatures to generate compositional shifts due to disproportionate attrition of endotherms and/or range expansion of ectotherms. If such trends are sustained, this will eventually lead to relatively higher occupancy, abundance, and richness of ectothermic vertebrates as pressure from endothermic predators and competitors declines.

To generate a simple forecast of how the relative balance of endotherms and ectotherms will change under different warming scenarios, we used CMIP 6 climate modeling scenarios for 2100 adopted by the Intergovernmental Panel on Climate Change^33,34^. These included an optimistic scenario of a global 1.4 °C increase (SSP SSP1–2.6)^35^, a more realistic 2.6 °C increase (SSP2–4.5)^36^, and a high-emissions, ‘business as usual’ scenario resulting in a 5.4 °C increase (SSP5–8.5)^37^. We adopted a ‘space for time substitution’ approach, which has been widely used to predict how species distributions, trait frequencies, and communities will shift as the climate changes^38,39^. We focus on temperature to model future diversity because of its central causal role, to define the scope of inference, and to evaluate the implications of our findings. To illustrate future global shifts in vertebrate diversity, we examine two comparable endothermic and ectothermic taxa — mammals and reptiles — which are primarily terrestrial, quadrupedal, and are not reproductively tied to water.

Today, mammals are more speciose than reptiles on 68% of the Earth’s surface where both coexist (Fig. 6A; mean ratio = 3.68, SD = 6.56). To forecast how this balance will shift, we used the observed relationship between richness and climate to adjust the baseline ratio based on future climate scenarios. Under a 1.4 °C global increase, the fraction of land with a mammalian majority declines to 54% (Fig. 6B); at 2.6 °C it falls to 46% (Fig. 6C). With a 5.4 °C increase, the pattern is nearly reversed: reptiles are more speciose on 65% of the Earth’s surface (Fig. 6D; mean ratio = 1.83, SD = 3.10). Notably, reptile diversity increases relatively more in drier areas. Indeed, two areas on the globe with the highest proportional reptilian diversity are the Sahara Desert and the arid interior of Australia (Fig. 6D). The relatively higher diversity of reptiles in deserts may reflect their lower metabolic demands, permitting greater occupancy in these unproductive deserts. These forecasts do not represent immediate or short-term shifts; rather, they are new diversity equilibria that vertebrate communities are expected to move toward under the sustained effects of climate change.

**Figure 6:**
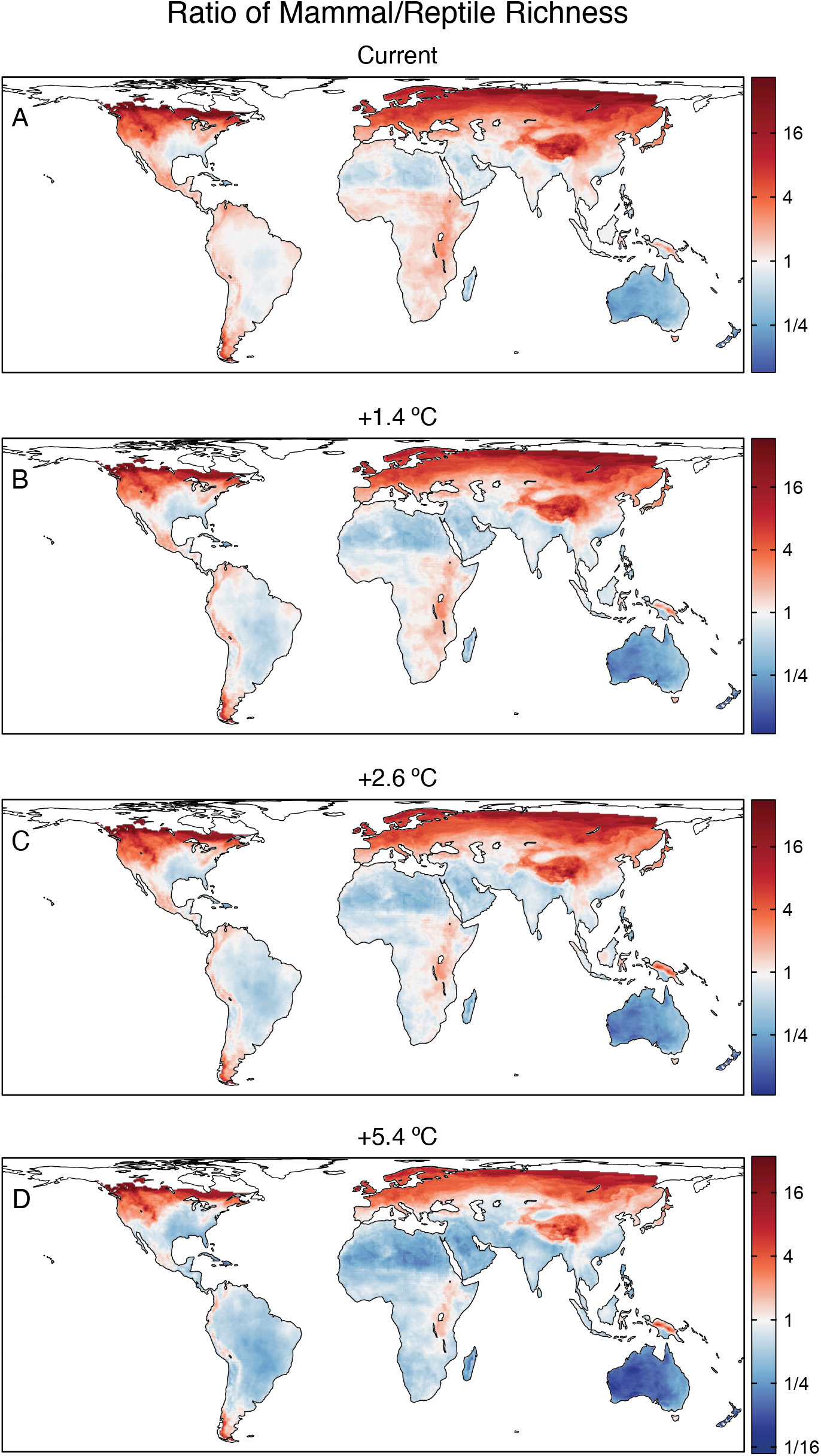
Comparative diversity of mammals and reptiles under different warming scenarios. In all plots, red indicates more mammal species compared to reptiles, blue indicates more reptile species than mammals, and white indicates an equal number. **(A)**. Currently, 68% of the Earth’s land surface area has more mammal species than reptile species. **(B-D)** Under multiple CMIP6 warming scenarios, the ratio of mammal to reptile species declines. Under the warmest scenario (D), the current pattern is reversed, with reptiles being more diverse across 65% of the Earth’s land surface (n cells = 50,879.

## Discussion

It has long been recognized that species diversity increases toward the tropics. Ambient temperature also increases toward the tropics and temperature appears to be the single best environmental predictor of richness for many taxa^3,40^. If this relationship is causal at a physiological level, thermoregulatory strategy — which determines internal body temperature — should be an equally strong predictor of vertebrate diversity. While there are other constraints on absolute richness, including habitat size and productivity, we hypothesized that the relative composition of vertebrate communities is primarily shaped by temperature. Supporting this prediction, the comparative richness of vertebrate endotherms and ectotherms varied systematically with ambient temperature (1, Fig. S3), while other variables offered little explanatory power. Strikingly, the magnitude of this shift is almost identical to the thermal sensitivity of metabolic rate, underscoring theoretical arguments that metabolism is a structuring force of global diversity^3,6,16^. The consequence is a global reorganization of vertebrate diversity with latitude and elevation: from endotherm dominance in cold ecosystems toward parity with ectotherms in the lowland tropics where body temperatures converge.

This striking pattern suggests a thermal mechanism — yet testing global theories of diversity is challenging. Metabolic frameworks, for instance, often rely on large-scale syntheses but rarely undergo experimental validation. Addressing this gap requires direct tests that connect individual thermal responses to broader diversity patterns.

Metabolic Asymmetry Theory (MAT) posits that differences in spatial diversity between endotherms and ectotherms reflect the temperature-dependence of species interactions^6^. Endothermic predators and competitors outperform ectotherms in colder environments due to their higher relative performance, leading to greater relative resource capture, abundance, and richness. While MAT has ecosystem-scale support^5,6^, it has lacked empirical validation at the individual level. We addressed this gap experimentally by quantifying key predatory metrics — such as time to capture and strikes per hunt — and showing that they reflect differences in thermal sensitivity of locomotion between predator and prey, as predicted by theory (Fig. 4). Our work also addresses a secondary issue: despite substantial learning and improvement in hunting behavior with experience, mice were unable to overcome fundamental kinetic constraints on prey capture. Ultimately, the exponential increase in red runner speed with rising temperatures posed formidable challenges that learning could not surmount (Fig. 5).

### Theoretical Drivers of Richness

Our proposed role of temperature in shaping global biodiversity shares important similarities and differences with other metabolic models of diversity. In their influential framework, Allen and colleagues proposed that metabolism drives global diversity patterns by regulating population size and evolutionary rates along thermal gradients^3,16,41^, leading to widespread temperature–richness slopes of *E* = 0.65 for ectotherms. While we focus on comparative shifts in endotherm and ectotherm diversity, these patterns can be analytically decomposed into the difference between the temperature-richness slopes of each group — the central focus of Allen et al. (2002).

Viewed through a complementary lens, our model offers a mechanism for why total community abundance of ectotherms may remain relatively constant with temperature — as observed by Allen et al. (2002) — despite declining per capita metabolic demands. Under an energy equivalence framework^42^, lower per capita metabolic rates in cold climates should lead to higher total ectothermic densities, since each individual needs fewer resources to exist. MAT offers a resolution — elevated predation and competition from endotherms constrain ectotherm densities in the cold.

Further, under our model, both the presence and absence of endothermic interactions help explain ectothermic convergence or divergence from an *E* = 0.65 slope of richness. In particular, we expect ectotherms that experience strong competition or predation from endotherms will be less diverse in cold habitats than those with weaker interactions, strengthening temperature–diversity associations. Supporting this prediction, squamates — which are terrestrial and conspicuously active during the day — exhibit a steeper thermal sensitivity of richness (*E* = 0.69), whereas cryptic, nocturnal amphibians and armored, aquatic turtles show much shallower slopes (*E* = 0.37 for frogs, *E* = 0.12 for turtles; Supp. Data, Tab 1). Indeed, temperature-richness slopes near *E* = 0.65 may depend on robust endotherm–ectotherm interactions, as lower slopes are often found in taxa that are too small-bodied to interact with endotherms. For instance, marine copepods exhibit a thermal sensitivity of richness around *E* ~ 0.3 (ref.^43^), while a continental analysis of soil bacteria reported a value near *E* ~ 0.2 (ref.^44^).

### Ecological Consequences of Thermal Asymmetry

Regardless of the ultimate causes of diversity, our model also departs from previous work by focusing on the ecological consequences of metabolic asymmetry for species interactions and community structure. For endotherms, the challenge of capturing fast-moving ectothermic prey in warm conditions should not only lead to decreased densities and niche opportunities for many mammals and birds, but also drive functional and compositional turnover for those that remain. Notably, the most speciose family of terrestrial, ectotherm-hunting mammals — invertivorous shrews (Soricidae, *>* 400 spp) — are highly diverse in the temperate northern hemisphere but absent from the lowland neotropics, where similar niches are filled by slower-metabolizing marsupials in the New World and a slow-metabolizing shrew subfamily in the Old World (Fig. S4A).

This metabolic asymmetry also shapes interactions in the opposite direction: in warmer habitats, endotherms are expected to be more vulnerable to predation by high-performance ectotherms. Supporting this prediction is recent work showing that reptile predation on tropical bird nestlings declines with elevation, whereas mammal predation increases^45^.

In contrast, ectotherms should be at greater risk of predation at higher latitudes and elevations, which may lead to range reduction, local extinction, and a reduction of realized niches. We suggest that this is especially true for large ectotherms, which have more limited opportunities to hide and avoid detection — an effective low-energy strategy that reduces their exposure to endothermic predators. For instance, catches of large predatory fish shift from open-water pelagic fish in the tropics to deeper demersal fish in temperate zones, where mammal and avian predators abound^46^. Predation risk may also help explain why endotherms tend to follow Bergmann’s rule in having larger body sizes in cold habitats,^47^ but reptiles do not^48^. In this case, selective pressures favoring larger body sizes in reptiles at high latitudes are counterbalanced by a greater vulnerability to endothermic predators.

The challenges ectotherms face from endotherms in cold climates may have a macroevolutionary signature. We anticipate that a latitudinal gradient of comparative diversity emerged soon after the advent of elevated body temperatures. In the late Cretaceous, for instance, warm-bodied dinosaurs have been found in cold arctic environments, but large ectotherms are absent^49^. In addition, the low diversity of large ectotherms today is a geological anachronism – out of step with much of earth’s history^50^. Giant, lumbering reptiles and synapsids once roamed Permian and Triassic landscapes free from endothermic predators, but it is unlikely that buffalo-sized iguanas, for instance, could survive the perils of warm-blooded lions today.

Interestingly, the observed latitudinal gradient between endotherms and ectotherms may help stabilize ecosystems at large spatial scales. In an ‘energy equivalence’ world^42^, higher metabolic rates at warmer temperatures should lead to lower ectotherm population densities and biomass, since each individual requires more resources. Before the evolution of endothermy, this would have produced a latitudinal gradient of vertebrate abundance declining toward the tropics. Today, however, vertebrate communities are characterized by warm ectotherms in the tropics and warm endotherms near the poles. Warm body temperatures and high metabolism at all latitudes help equalize per capita energy use, stabilizing vertebrate biomass and population densities across the globe.

### The Future of Vertebrate Diversity

As global temperatures rise, we forecast that relative richness will increasingly trend toward ectotherms. Because richness reflects longer macroevolutionary processes like speciation, large shifts in species numbers may not be immediately observed, but instead represent an equilibrium state that ecological communities will gradually approach. In the short term, changes in relative abundance will likely occur first. Next, range expansion of tropical species into temperate zones will occur, but thermally sensitive ectotherms should benefit the most. Some endothermic species will suffer, especially those feeding on agile ectothermic prey, such as small, invertivorous mammals and birds on land, and piscivorous penguins and pinnipeds at sea. Less impacted will be cognitively flexible predators like dolphins, and groups unaffected by metabolic asymmetry, such as endothermic herbivores and their warm-bodied predators. Nonetheless, a broad, proportional decline in the composition and abundance of endotherms in a warming world will open up more opportunities for reptiles and fish. Climate change imposes constraints on all animals but the effects of hotter temperatures will not be distributed equally. We expect the future of life on land to become increasingly reptilian.

## Acknowledgments

This project was supported by the Living Earth Collaborative at Washington University in St. Louis (J.M.G.). J.M.G. and A.I.D. were supported by a National Science Foundation grant DEB-1838346. Support was also provided by the National Institutes of Health BRAIN (Brain Research through Advancing Innovative Neurotechnologies) Initiative award R01NS118442 (K.B.H.) and the National Institutes of Health, Institutional national research service awards: T32EY013360 from the National Eye Institute and T32NS121881 from the National Institute of Neurological Disorders and Stroke (J.L.A). We thank Michael Moore, David van Horn, James Stroud, and Andrea L. Martin for helpful suggestions, and Marcos Bermudez and Nicole Paquette for assistance in running experiments.

## Author Contributions

J.M.G. and A.I.D. developed theory; J.M.G., A.I.D., J.L.A. and K.B.H. designed the experiments. J.M.G. and J.L.A. performed experiments. J.M.G., J.L.A., K.B.N., S.J.B., Q.D.R., and V.S. analyzed data. J.M.G., J.L.A., and K.B.H. wrote the manuscript. A.I.D. and K.B.H. supervised the project. All authors interpreted the results and reviewed and revised the manuscript.

## Data Availability

All spatial data is publicly available (see Methods); experimental data is available upon request.

## Code Availability

All code for analyses are available at https://github.com/hengenlab/temp_dependent_predation.

## Methods

### Terrestrial Vertebrate Distributions and Diversity

Species distributions for vertebrates were acquired from publicly available datasets and publications. Mammal and amphibian distributions were obtained from the IUCN^19^, bird distributions from BirdLife International^51^, and reptile distributions from Roll et al. (2017)^52^. Elevation, annual precipitation and annual surface temperature data were obtained from WorldClim 2^53^, and NPP (net primary productivity) data from NASA Earth Observations^54^.

To calculate species richness, we used the Behrmann equal area projection with a WGS84 grid of 48.25 km X 48.25 km, corresponding to ~ 0.5° X 0.5° at 30° latitude. A species occurrence was recorded wherever a species distribution overlapped any part of the grid; vagrants and non-native species were excluded. Occurrences were then summed to calculate richness. Only terrestrial areas and species were included (see Grady et al. (2019) for a list of marine tetrapods that were excluded^6^). We also performed a sensitivity analysis on spatial grain size, assessing diversity patterns at 2X and 4X resolution. The thermal sensitivity of endotherm/ectotherm richness was insensitive to grain size to two decimal points (linear model: *E* = 0.58 for all resolutions).

Global analyses of vertebrate diversity tend to focus on a few environmental drivers, including temperature, elevation, net primary productivity (NPP), and precipitation^2^. To calculate the change in richness with temperature, we used the following formulation: log(endo rich/ecto rich) ~ *E* (1/kT) + log(precipitation) + log(NPP) + elevation range, where *k* is Boltzmann’s constant (8.617 eV per Kelvin), *T* is temperature in Kelvin, *E* is the slope and thermal sensitivity coefficient (*E* ~ 0.65 for metabolic rate^18^). These predictor variables had a low variance inflation factor (*<* 3), indicating low multi-collinearity (package *car* ^55^). Substitution of elevational standard deviation for elevational range did not change results. Of these four predictors, temperature was the most important factor by far. Partial *r*^2^ values for temperature = 0.75, precipitation = 0.0060, and elevation = 0.0097, NPP = 0.00036; linear model analyzed using the package rsq^56^).

We accounted for spatial autocorrelation in our analysis by using the Besag-York-Mollie 2 (BYM2) model^57^, implemented in R-INLA^58^. BYM2 is a spatial Hierarchical Bayesian model that includes fixed effects (environmental predictors) and random effects, partitioning random effects into spatially structured and unstructured components. It uses penalized complexity priors to avoid overfitting. R-INLA employs efficient integrated nested Laplace approximations of posterior distributions. Spatial adjacency was defined using a distance-based neighborhood matrix, with cells considered neighbors if within 1.5 times the grid resolution. Penalized complexity (PC) priors were used for the spatial random effect following the BYM2 formulation^59^. The prior on the marginal standard deviation was set as P(SD *>* 1) = 0.01, and the prior on the proportion of spatially structured variance (*ϕ*) was set as 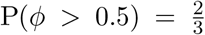. These priors favor simpler models while allowing moderate spatial structure when supported by the data. INLA approximates posterior distributions deterministically, without requiring Markov chain Monte Carlo (MCMC) sampling. Results from this model (shown in Fig. 2, see also Fig. S3) are qualitatively similar to multiple linear regression results (see Supp. Data 1, Tabs 1-2). In addition, *E* for richness was robust to the exclusion of non-thermal predictor variables (full BYM2 model: *E* = 0.64, *r*^2^(*fixed*) = 0.79; BYM2 temperature only: *E* = 0.60, *r*^2^(*fixed*) = 0.61; full linear model *E* = 0.58, *r*^2^ = 0.80, linear model, temperature only: *E* = 0.55, *r*^2^ = 0.77). n spatial cells for all models is 54,795.

We use log-transformed ratios of richness(e.g., log(Endotherm:Ectotherm) as response variables in our analysis because they offer interpretive, mechanistic, and statistical advantages. First, they match our *comparative focus*: rather than modeling each group’s richness independently, the ratio directly captures proportional diversity and dominance — where over space one group outnumbers another. This aligns with both the goals of our analysis and our theoretical framing: species interactions are inherently comparative, and differences in richness emerge through relative dominance. This logic parallels other fields where relative performance is key: economics (e.g., debt-to-GDP), microbiome science (e.g., Firmicutes:Bacteroidetes), and phylogenetics (e.g., lineage-through-time ratios). Second, ratios offer *mechanistic insight* : environmental stressors like freezing or drought often reduce richness across taxa, creating strong nonlinearities in absolute richness with latitude or temperature. Ratios help offset this shared signal and isolate shifts in balance due to physiology or interaction strength. Finally, log ratios have *statistical advantages*: in our case, they reduce skew and heavy tails, produce more normal residuals, and improve linearity (Data 1: Tab 3) without sacrificing fit (Data 1: Tab 1-2). And because the log of a ratio is symmetric `(log(A/B) = {log(B/A))`, the directionality of dominance does not bias the analysis — it only flips the sign.

### Climate Projections

Data for temperature forecasts was obtained from Intergovernmental Panel on Climate Change Sixth Assessment projections, based on climate scenarios known as Shared Socioeconomic Pathways (SSPs)^34^. We used output from the CESM2 model within the Coupled Model Intercomparison Project Phase 6 (CMIP6)^33^. Specifically, scenario headers in figures report projected global average temperature increases by 2100 relative to the year 2014, to indicate how temperatures will warm from the present. This incldes 1.4 °C for SSP1-2.6 (low emissions)^35^, 2.6°C for SSP2-4.5 (‘Middle of the Road’)^36^, and 5.4°C for SSP5-8.5 (high emissions)^37^. CESM2 climate data was downloaded from https://cds.climate.copernicus.eu, and reprojected to 48.25 km x 48.25 km Behrmann equal area grids.

For all modeled projections, we used climate anomalies relative to the 1970–2000 mean—the period covered by the WorldClim observational climate dataset that served as the baseline for all spatial regression model fitting and richness projections. As a result, there is a small offset between the scenario temperature increases shown in figure headers (relative to 2014) and the baseline used for projections (1970–2000). This approach provides readers with scenario values that are easily interpreted as ‘warming above present-day,’ while maintaining methodological consistency in spatial modeling.

To forecast changes in comparative richness (Fig. 6), we first used current comparative richness of mammal to reptiles as a baseline (see Fig. 6A). Next, we calculated the linear regression model fit between mammal/reptile richness and used fitted coefficients for temperature and precipitation for predictions: lm(log(mammal rich/reptile rich) ~ (temperature C) + log(precipitation) + (elevation range): log(mammal richness/reptile richness) = −0.24 − 0.084(temperature) + 0.11log(annual precipitation); all variables: *p <* 0.001). We then calculated the deviations in temperature and precipitation between current conditions and climate model projections for all spatial cells. These climate deviation were multiplied by the regression fit, yielding projected deviations of diversity. Finally, we added projected deviations to current endotherm/ectotherm richness to generate future estimates of comparative diversity. We based forecasts on linear models rather than BYM2 models with spatial autocorrelation in order to reflect empirical autocorrelation relevant to future diversity (i.e., the richness of a given cell is indeed influenced by neighboring cells).

### Metabolic Asymmetry Theory

In ectothermic animals, metabolic and locomotory rates generally increase with temperature until reaching a peak and then decline. The thermal dependence of metabolic rate and locomotory rates *R* during the upward portion of the curve is described by Eq. 1: *R* ∝ *e*^*−E/kT*^, where *k* is Boltzmann’s constant, *T* is temperature in Kelvin and *E* is a thermal sensitivity coefficient (‘activation energy’), which is ~ 0.65 for metabolic rate^18^ and ~ 0.5 for speed^30^. In our experiments, red runner speed increased until reaching a plateau at 30 °C. Therefore, we calculated *E*_*Speed*_ from 14 °C to 30 °C. For comparison, we only report estimates of *E*_*Speed*_ in mice up 30 °C.

Previously, we published a theoretical model — Metabolic Asymmetry Theory (MAT) — to explain elevated marine endotherm diversity at high latitudes^6^. We proposed that the asymmetric metabolic and performance responses to ambient temperature between endotherms and ectotherms give endotherms a performance advantage in cold environments, as ectothermic prey, predators, and competitors become cold and sluggish. At an ecosystem scale, we found support for MAT, showing that that the fraction of prey captured by endothermic predators relative to ectothermic predators was a function of temperature:

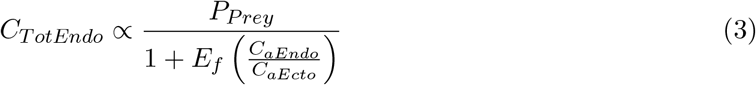

where, *C*_*Tot*_ is total consumption, *C*_*a*_ is maximum individual consumption, and *P*_*Prey*_ is prey productivity. The thermal sensitivity of global marine mammal consumption *E*_*f*_ fell within the predicted range^6^.

A key prediction of MAT at the individual scale is that capture efficiency *C*_*E*_ is proportional to the ratio of predator to prey speed: *C*_*e*_ ∝ *S*_*Pred*_*/S*_*Prey*_, where *C*_*e*_ is defined as the capture per encounter in a motivated predator. Equivalently, escape efficiency *ϵ* is the inverse (Eq. 2: *ϵ* ∝ *S*_*Prey*_*/S*_*Pred*_ ∝ Δ*E*^*α*^, where Δ*E* is the difference in thermal sensitivity of speed between predator and prey (*E*_*SPrey*_ − *E*_*SPred*_) and *α* ≤ 1. If endotherm performance is thermally invariant, *E*_*SPred*_ = 0 and Δ*E* simplifies to *e*^*−E/kT*^. For predators pursuing fast-moving prey, *α* is expected to be ~ 1, but it may be lower under two conditions. First, if prey are much slower, *α* ~ 0. That is, as the *S*_*Prey*_*/S*_*Pred*_ → ∞, *α* → 0. For instance, thermal changes should have little effect on escape efficiency when endotherms forage for plants or snails. Second, *α* may be less than 1 if predators use cooperative behavior to neutralize prey speed. This appears to be the case with tropical dolphins, which herd fish into churning ‘bait balls’ at the water’s surface^60^, or humans using nets.

In our experiments, capturing red runners required active pursuit and was relatively challenging, especially at warmer temperatures. Additionally, mice are not cooperative hunters, so *α* = 1, and Δ*E*^*α*^ simplifies to Δ*E*. Because red runners were in a confined arena, capture is assured so long as predators are motivated. Thus, we used time to capture as a proxy for escape efficiency. In nature, longer times to capture should correspond to a higher efficiency of escape.

In our experiments, Δ*E* was predicted to be 0.5 −0.65, with the lower bound matching observed thermal sensitivity of ectotherms speed in empirical syntheses^30^, and the upper bound reflects the thermal sensitivity of respiration in macroecolgoical analyses^18^. We focus on Δ*E* evaluated from animals in isolation to measure innate differences in performance capacity relevant to capture success. Further, we focused on differences in the thermal sensitivity of speed rather than acceleration, as capture involved unobstructed pursuit around the arena’s perimeter (for ambush predators acceleration may be more relevant). We used the difference in thermal sensitivity of median and maximum speed between predator and prey in isolation to establish a range of predicted *E*_*CaptureTime*_ (Fig. 4G).

### Experimental model and subject details

All experimental procedures were conducted in accordance with protocols approved by the Washington University in Saint Louis Institutional Animal Care and Use Committee (IACUC), following NIH guidelines for the care and use of research animals. Five female and three male C57BL/6J mice (Supplier: The Jackson Laboratory) were used in this study. Following initial trials at room temperature, one male mouse stopped hunting and was excluded from subsequent thermal trials (see below for experimental details). Mice were seventeen weeks old at the time of prey capture experiments. We observed no differences in predation between male and female mice and, therefore, combined their data. Turkestan red runner cockroaches (Supplier: Caribbean Mealworms) aged three to four months and weighing 0.05 - 0.20 g (mean = 0.11 g), were used as prey for all capture experiments.

### Prey Capture Experiments

Forty-eight hours prior to the commencement of hunting trials, mice were housed individually in circular hunting arenas (36.83 cm H x 25.87 cm D) and provided with food pellets and water ad libitum. The arenas, which also served as home cages for the duration of the experiments, were located in ventilated and temperature-controlled chambers on a 12:12 light/dark cycle (Fig. S5E-G). After forty-eight hours of the mice habituating to these new enclosures, three red runners were introduced to each room temperature (i.e., 25 °C) arena overnight (i.e., 12 h dark period), alongside ad libitum access to food pellets. Any red runners that remained at the end of the 12 h dark period were then removed. This protocol was repeated for 1 d for a total of 2 d of nocturnal exposure to red runners.

We then conducted ten days of initial hunting trials at room temperature (Fig. 5, left column; Fig. S6). Each trial consisted of a red runner being introduced to the arena, and the interactions of mice and red runners were recorded with an overhead camera (30 fps; e3Vision; White Matter). Each mouse had the opportunity to capture up to six red runners daily (~ 60 trials total per mouse), with a ten minute delay between trials (Fig. S5D). Trials were completed when the red runner was incapacitated. Food pellets were removed 14 - 16 h before each day of hunting trials, which commenced 1 h after the onset of the light period. After the conclusion of each day of hunting trials, food pellets were returned to the arenas for 8 - 10 h (Fig. S5D).

After the initial trials, we conducted thermal trials at 14, 18, 25 (room temperature), 30, and 35 °C (Fig. S5F; 5, right column; Fig. S10). Protocols were otherwise the same as initial trials. Both red runners and mice were kept in a shared temperature-controlled chamber, ensuring they experienced the same thermal conditions. Thermal trials were conducted in two experimental phases. In the first phase, which was conducted for 27 d, the target temperature was switched every 5 d (~ 30 trials total per temperature per mouse). To control for thermal trial order effects, half of the mice experienced cold conditions before warm and half experienced warm conditions before cold. To check if temperature-dependent performance was robust after weeks of training, we also conducted a second phase of thermal trials for 10 d, in which the target temperature was switched every 1 d. We observed no differences in predation between phases; therefore, we pooled all data for analyses.

Trials were excluded if mice showed signs of overheating at 35 C (e.g., splaying legs), were not persistent in hunting, failed to eat prey, or recording errors occurred, totaling 13.1% of trials.

### Thermal Mismatch Trials

Following thermal trials, we conducted thermal ‘mismatch’ trials to test whether mice use thermal cues to anticipate prey behavior. Standard thermal trials are matched — the red runner has the same internal body temperature as the ambient hunting arena. Thus, mice can potentially learn to associate ambient temperature with prey behavior: e.g., fast-moving prey in warm arenas and slower prey in cold arenas. To test this possibility, we conducted mismatch trials. In a mismatch trial, mice in cold arenas (14 °C) that normally receive cold red runners occasionally received a warm red runner (30 °C), and vice versa. We then compared the locomotion and hunting performance of mice in these mismatched conditions to their performance in standard, thermally matched conditions. In particular, the performance of mice hunting warm red runners in warm arenas (‘standard warm’) was compared to the performance of mice hunting warm red runners in cold arenas (‘mismatch warm’), and likewise for hunting cold red runners in ‘standard cold’ and ‘mismatch cold’ conditions (Fig. S11A,C).

Thermal mismatch experiments were conducted for 12 days total. Half of mice started trials in cold conditions (14 °C); the other half in warm conditions (30 °C). After 6 days, the conditions were switched. To maintain potentially learned associations between ambient temperature and prey behavior, most trials followed standard protocol: across each set of 6 days, matched thermal trials were carried out as previously described, with 6 trials per mouse per day. Every other day a thermal mismatch trial was performed, followed by five matched thermal trials. Thus, over 12 days, 6 mismatch trials occurred for each mouse (for seven mice), for a total of 42 mismatch trials. For statistical comparisons, we only considered the first trial per day: 42 trials under mismatched conditions and 42 trials under standard conditions. Although thermally mismatched red runners eventually warmed or cooled toward ambient temperature, they were typically consumed within 3 to 20 s (Fig. S11B,D), limiting the change in red runner body temperature (Fig. S12).

Effect sizes were calculated using Cohen’s *d*, defined as the model-adjusted difference in marginal means divided by the pooled standard deviation of model residuals. Confidence intervals for *d* were estimated using a normal approximation, with standard error derived from group sample sizes and the effect magnitude. See Data 1, Tab 8 for results.

### Movement Tracking and Analysis

Videos of mice and red runners were recorded with an overhead camera at 30 frames per second (e3Vision, White Matter). Each trial was manually scored for time to capture and scoring accuracy was verified on a subset of videos by two trained scorers. Markerless pose estimation was conducted on the mouse and red runner using DeepLabCut^61^ (Fig. 3A), specifically tracking six anatomical points on the mouse (snout, left ear, right ear, shoulder, spine, tail base) and two points on the red runner (front and back). Based on these six anatomical points, other points were extracted using midpoint calculation 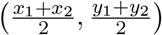, e.g., red runner center. After this, second-order features such as distance traveled 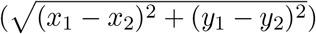 velocity (| 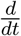distance traveled| × fps), and acceleration (| 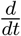velocity|) were calculated for each animal (custom Python script). Strikes between mouse and red runner were defined as when both points on the red runner were occluded by the mouse for 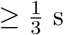 s (Fig. 3C). Red runner movement was calculated with respect to red runner center while mouse movement was calculated with respect to mouse spine.

### Video Post-processing

Markerless pose estimation was conducted on the mouse and red runner videos using DeepLabCut^61^ across 9 million total frames. Occasionally, the DeepLabCut model would produce erroneous pose estimate values, so a series of post-processing steps was conducted to identify and remove outliers. First, for each video, a frequency distribution of the change in spatial position from one frame to the next was inspected for outliers and then compared to the corresponding video segment to determine a conservative threshold for exclusion. Specifically, pose estimate values were excluded if the distance traveled from one frame to the next exceeded 3 cm (equivalent to 90 cm/s). Additionally, red runner pose estimate values were discarded if the front and back values differed by more than 2 cm (i.e., exceeding body length).

Next, we sought to exclude recorded movement values that did not reflect actual locomotion; e.g., grooming or pixel jittering. We first assessed videos of animals visually confirmed to be non-locomotory (n = 63 for red runners, n = 60 for mice). The 95% quantile of these non-locomotory values was treated as a threshold separating non-locomotory and locomotory movement; values below this threshold were subsequently excluded in analyses (Fig. S9).

Finally, we used the following formula to identify and exclude remaining outliers: *Q*_2_ *±C* ×IQR, where *Q*2 is the median, *IQR* is the 25 − 75% inter-quartile range, and *C* is a coefficient (note that *C* = 1.5 is the outlier range in a traditional box plot, equivalent to 2.7 standard deviations in a normally distributed dataset). For a given trial, we used a conservative value of *C* = 3 (equivalent to 4.5 standard deviations), and for pooled red runners or mice at each thermal condition, we used an even more conservative *C* = 6 (equivalent to 9 standard deviations).

### Temperature Recordings

To measure mouse body temperature, mice were placed in temperature-controlled chambers (14 °C, n = 4; 30 °C, n = 4) for 30 minutes and then rapidly anesthetized with 5% isoflurane. Upon confirmation of loss of consciousness (~ 10 s), each mouse was decapitated and then a sterile temperature probe was immediately inserted into the brainstem via the foramen magnum. Temperature data was collected for 5 s at a frequency of 1 Hz and were recorded with a Leaton Digital Dual Thermometer with K Type Thermocouple (Fig. S7A).

To measure roach internal temperature, red runners were placed in temperature-controlled chambers (14 °C, n = 4; 18 °C, n = 2; 25 °C, n = 4; 30 °C, n = 11) for 30 minutes and then a sterile temperature probe was inserted into the abdomen of each red runner. Temperature data was collected for 5 s at a frequency of 1 hz and were recorded with a Leaton Digital Dual Thermometer with K Type Thermocouple (Fig. S7B).

For thermal imaging, naive mice and red runners were placed in temperature-controlled chambers that were either cooled or heated to 14 °C or 30 °C, respectively. After 30 minutes, one mouse and one red runner from matched thermal conditions (i.e., 14 °C and 30 °C) were paired together and imaged using a thermal camera (FLIR E53; Teledyne FLIR). This imaging protocol was repeated with mice and red runners from mismatched thermal conditions (Fig. S11A,C).

### Statistical Analysis of Experiments

All statistical analyses were performed in R, using the ‘tidyverse’ package for data processing and visualization. Repeated measures of mice were addressed with mixed model regression, using packages and ‘lme4’, ‘lmerTest’, and ‘emmeans’ packages. Nonlinear fits in Fig. 3 and Fig. S6 are LOESS fits generated in ‘ggplot2’. See code for full details.

A linear mixed model was fit to log-transformed response rates to estimate the thermal sensitivity parameter *E*:

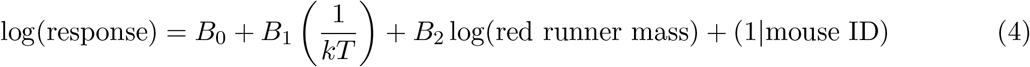

where *B*_0_ is the intercept, *B*_1_ is the thermal coefficient and estimate of *E, B*_2_ is the coefficient associated with red runner mass, *k* is Boltzmann’s constant ( 8.617 × 10^*−*5^ eV/K), *T* is temperature in Kelvin, and mouse ID is a random effect. Red runner mass was included for its predictive value (Fig. S8), and theoretical significance, as large animals tend to be faster^62^ and may be more challenging to overpower.

We evaluated model assumptions using residual diagnostics, including Q–Q plots and quantitative assessments of normality. Specifically, we calculated skewness, excess kurtosis, and the root mean square error (RMSE) of the standardized residuals against a normal Q–Q line. Residuals were approximately normal within ±2 standard deviations, with moderate tail deviations in some cases. While formal normality tests are overly sensitive given the large sample size and spatial autocorrelation, these metrics indicated no meaningful departures from normality that would compromise inference. It is also important to note that regression of the ratio of endotherm: ectototherm richness was more linear than regression of either alone, with lower skew, kurtosis and QQ RMSE (Data 1, Tab 3).

In addition, we used corrected Akaike Information Criterion (AICc) to analyze the role of the following potential predictor variables as main effects: mouse mass, mouse sex, trial number, day of trial, and order of heating/cooling trials. We also considered the following random effects: experimenter ID, time-to-capture scorer ID, and mouse ID. We performed model comparisons using the MuMIn::AICc() and lmerTest::step() functions in R, with the latter applying backward selection based on AIC values. For all response variables, temperature, red runner mass, and mouse ID consistently yielded the lowest AIC values. Occasionally, for specific response variables, mouse mass and trial number contributed to models with the lowest AIC values. However, estimates of the primary parameter of interest — temperature — remained insensitive to inclusion or exclusion of additional predictors. To maintain parsimony and facilitate interpretation, reported results focus solely on the variables shown in *Eq. 4*.

For comparisons in Figure 5, we used linear mixed-effects models to assess the effects of temperature (‘hot’ vs ‘cold’ conditions) on behavioral and locomotor response variables in mice, while accounting for repeated measures from individual animals. Hot refers to trials conducted at 30 °C or 35 °C and cold is 14 °C or 18 °C. For each response variable, a mixed model was fitted with temperature regime, trial sequence, and covariates (such as prey mass) as fixed effects, and mouse ID as a random effect to control for within-subject correlation.

Model-based marginal means for each temperature condition were estimated using the emmeans package. The difference between hot and cold conditions was evaluated using model-derived pairwise contrasts. For each variable, we report the estimated mean difference (on the log scale), 95% confidence interval, test statistic, degrees of freedom (rounded to the nearest integer), p-value, and sample size for each group. See Data 1, Tab 7 for results.

**Supplemental Data 1. Statistical results for spatial and experimental analyses. Tab 1** presents statistical outputs for Bayesian models of temperature and vertebrate diversity, which account for spatial autocorrelation. Predictor variables include ambient temperature, precipitation, net primary productivity (NPP), elevation range, as well as random spatial effects. Temperature was also considered as the only fixed effect. **Tab 2** presents similar results but for a linear model, excluding random effects. **Tab 3** presents residual normality analyses for models of species richness, evaluated separately for endotherms, ectotherms, and their richness ratio (endotherms : ectotherms). **Tab 4** shows shrew metabolic rates taken from the literature (citations included). **Tab 5** provides statistical results for experimental analysis of acceleration with temperature. **Tab 6** shows results from initial experimental trials at room temperature. **Tab 7** shows results from thermal trials. **Tab 8** shows results from thermal mismatch trials.

## References

1. Hillebrand, H. On the generality of the latitudinal diversity gradient. The American Naturalist 163, 192–211 (2004).

2. Raz, T. et al. Diversity gradients of terrestrial vertebrates–substantial variations about a common theme. Journal of Zoology 322, 126–140 (2024).

3. Allen, A. P., Brown, J. H. & Gillooly, J. F. Global Biodiversity, biochemical Kinetics, and the energetic-equivalence rule. Science 297, 1545–1548. issn: 1095-9203. 10.1126/science.1072380 (Aug. 2002).

4. Buckley, L. B., Hurlbert, A. H. & Jetz, W. Broad-scale ecological implications of ectothermy and endothermy in changing environments. Global Ecology and Biogeography 21, 873–885 (2012).

5. Cairns, D., Gaston, A. & Huettmann, F. Endothermy, ectothermy and the global structure of marine vertebrate communities. Marine Ecology Progress Series 356, 239–250. issn: 1616-1599. 10.3354/meps07286 (Mar. 2008).

6. Grady, J. M. et al. Metabolic asymmetry and the global diversity of marine predators. Science 363. issn: 1095-9203. 10.1126/science.aat4220 (Jan. 2019).

7. Santini, L. et al. Global drivers of population density in terrestrial vertebrates. Global Ecology and Biogeography 27, 968–979 (2018).

8. Roslin, T. et al. Higher predation risk for insect prey at low latitudes and elevations. Science 356, 742–744. issn: 1095-9203. 10.1126/science.aaj1631 (May 2017).

9. Caley, M. J. & Schluter, D. The relationship between local and regional diversity. Ecology 78, 70–80 (1997).

10. Arita, H. T. & Rodríguez, P. Local-regional relationships and the geographical distribution of species. Global Ecology and Biogeography 13, 15–21 (2004).

11. Belmaker, J. & Jetz, W. Regional pools and environmental controls of vertebrate richness. The American Naturalist 179, 512–523 (2012).

12. Kissling, W. D., Sekercioglu, C. H. & Jetz, W. Bird dietary guild richness across latitudes, environments and biogeographic regions. Global Ecology and Biogeography 21, 328–340 (2012).

13. Kissling, W. D. et al. Establishing macroecological trait datasets: digitalization, extrapolation, and validation of diet preferences in terrestrial mammals worldwide. Ecology and Evolution 4, 2913–2930 (2014).

14. Willig, M. R., Kaufman, D. M. & Stevens, R. D. Latitudinal gradients of biodiversity: pattern, process, scale, and synthesis. Annual review of ecology, evolution, and systematics 34, 273–309 (2003).

15. Gillooly, J. F. & Allen, A. P. Linking global patterns in biodiversity to evolutionary dynamics using metabolic theory. Ecology 88, 1890–1894 (2007).

16. Stegen, J. C., Enquist, B. J. & Ferriere, R. Advancing the metabolic theory of biodiversity. Ecology letters 12, 1001–1015 (2009).

17. Storch, D. in Metabolic ecology: a scaling approach 120–131 (Wiley Online Library, 2012).

18. Gillooly, J. F., Brown, J. H., West, G. B., Savage, V. M. & Charnov, E. L. Effects of Size and Temperature on Metabolic Rate. Science 293, 2248–2251. issn: 1095-9203. 10.1126/science.1061967 (Sept. 2001).

19. IUCN. 2021.

20. McNab, B. K. Food habits, energetics, and the reproduction of marsupials. Journal of Zoology 208, 595–614 (1986).

21. Vogel, P. in Comparative physiology: Primitive mammals (eds Schmidt-Nielsen, K., Bolis, L. & Taylor, C. R.) 170–180 (Cambridge University Press, 1980).

22. Hurlbert, A. H., Ballantyne, F. & Powell, S. Shaking a leg and hot to trot: the e!ects of body size and temperature on running speed in ants. Ecological Entomology (2008).

23. Burnside, W. R., Erhardt, E. B., Hammond, S. T. & Brown, J. H. Rates of biotic interactions scale predictably with temperature despite variation. Oikos 123, 1449–1456 (2014).

24. O’Connor, M. P. et al. Reconsidering the mechanistic basis of the metabolic theory of ecology. Oikos 116, 1058–1072 (2007).

25. Landry Jr, S. O. The Rodentia as omnivores. The Quarterly Review of Biology 45, 351–372 (1970).

26. Butler, K. Predatory behavior in laboratory mice: Strain and sex comparisons. Journal of Comparative and Physiological Psychology 85, 243–249. issn: 0021-9940. 10.1037/h0035008 (1973).

27. Hoy, J. L., Yavorska, I., Wehr, M. & Niell, C. M. Vision Drives Accurate Approach Behavior during Prey Capture in Laboratory Mice. Current Biology 26, 3046–3052. issn: 0960-9822. 10.1016/j.cub.2016.09.009 (Nov. 2016).

28. Hoy, J. L., Bishop, H. I. & Niell, C. M. Defined Cell Types in Superior Colliculus Make Distinct Contributions to Prey Capture Behavior in the Mouse. Current Biology 29, 4130–4138.e5. issn: 0960-9822. 10.1016/j.cub.2019.10.017 (Dec. 2019).

29. Johnson, K. P. et al. Cell-type-specific binocular vision guides predation in mice. Neuron 109, 1527–1539.e4. issn: 0896-6273. 10.1016/j.neuron.2021.03.010 (May 2021).

30. Dell, A. I., Pawar, S. & Savage, V. M. Systematic variation in the temperature dependence of physiological and ecological traits. Proceedings of the National Academy of Sciences 108, 10591–10596. issn: 1091-6490. 10.1073/pnas.1015178108 (May 2011).

31. Bryce, C. M., Wilmers, C. C. & Williams, T. M. Energetics and evasion dynamics of large predators and prey: pumas vs. hounds. PeerJ 5, e3701 (2017).

32. Hughes, R. N., Kaiser, M. J., Mackney, P. & Warburton, K. Optimizing foraging behaviour through learning. Journal of Fish Biology 41, 77–91 (1992).

33. Eyring, V. et al. Overview of the Coupled Model Intercomparison Project Phase 6 (CMIP6) experimental design and organization. Geoscientific Model Development 9, 1937–1958 (2016).

34. Riahi, K. et al. The Shared Socioeconomic Pathways and their energy, land use, and greenhouse gas emissions implications: An overview. Global environmental change 42, 153–168 (2017).

35. Danabasoglu, G. NCAR CESM2 model output prepared for CMIP6 ScenarioMIP ssp126 2019. 10.22033/ESGF/CMIP6.7746.

36. Danabasoglu, G. NCAR CESM2 model output prepared for CMIP6 ScenarioMIP ssp245 2019. 10.22033/ESGF/CMIP6.7748.

37. Danabasoglu, G. NCAR CESM2 model output prepared for CMIP6 ScenarioMIP ssp585 2019. 10.22033/ESGF/CMIP6.7768.

38. Blois, J. L., Williams, J. W., Fitzpatrick, M. C., Jackson, S. T. & Ferrier, S. Space can substitute for time in predicting climate-change e!ects on biodiversity. Proceedings of the national academy of sciences 110, 9374–9379 (2013).

39. Lovell, R. S., Collins, S., Martin, S. H., Pigot, A. L. & Phillimore, A. B. Space-for-time substitutions in climate change ecology and evolution. Biological Reviews 98, 2243–2270 (2023).

40. Tittensor, D. P. et al. Global patterns and predictors of marine biodiversity across taxa. Nature 466, 1098–1101 (2010).

41. Brown, J. H., Gillooly, J. F., Allen, A. P., Savage, V. M. & West, G. B. Toward a Metabolic Theory of Ecology. Ecology 85, 1771–1789. issn: 0012-9658. 10.1890/03-9000 (July 2004).

42. Damuth, J. Population density and body size in mammals. Nature 290, 699–700 (1981).

43. Rombouts, I., Beaugrand, G., Ibaněz, F., Chiba, S. & Legendre, L. Marine copepod diversity patterns and the metabolic theory of ecology. Oecologia 166, 349–355 (2011).

44. Luan, L. et al. Integrating pH into the metabolic theory of ecology to predict bacterial diversity in soil. Proceedings of the National Academy of Sciences 120, e2207832120 (2023).

45. Londoño, G. A., Gomez, J. P., Sánchez-Martínez, M. A., Levey, D. J. & Robinson, S. K. Changing patterns of nest predation and predator communities along a tropical elevation gradient. Ecology Letters 26, 609–620 (2023).

46. Van Denderen, P. D., Lindegren, M., MacKenzie, B. R., Watson, R. A. & Andersen, K. H. Global patterns in marine predatory fish. Nature ecology & evolution 2, 65–70 (2018).

47. He, J., Tu, J., Yu, J. & Jiang, H. A global assessment of Bergmann’s rule in mammals and birds. Global Change Biology 29, 5199–5210 (2023).

48. Pincheira-Donoso, D. & Meiri, S. An intercontinental analysis of climate-driven body size clines in reptiles: no support for patterns, no signals of processes. Evolutionary Biology 40, 562–578 (2013).

49. Druckenmiller, P. S., Erickson, G. M., Brinkman, D., Brown, C. M. & Eberle, J. J. Nesting at extreme polar latitudes by non-avian dinosaurs. Current Biology 31, 3469–3478 (2021).

50. Smith, F. A. et al. Body size evolution across the Geozoic. Annual Review of Earth and Planetary Sciences 44, 523–553 (2016).

51. Bird species distribution maps of the world 2017.

52. Roll, U. et al. The global distribution of tetrapods reveals a need for targeted reptile conservation. Nature ecology & evolution 1, 1677–1682 (2017).

53. Fick, S. E. & Hijmans, R. J. WorldClim 2: new 1-km spatial resolution climate surfaces for global land areas. International journal of climatology 37, 4302–4315 (2017).

54. Center, N. G. S. F. Net Primary Production (NPP) https://neo.gsfc.nasa.gov/view.php?datasetId=MOD17A2_M_PSN. NASA Earth Observations (NEO) dataset, accessed on 2024-08-06.

55. Fox, J. & Weisberg, S. An R companion to applied regression (Sage publications, 2018).

56. Zhang, D. rsq: R-Squared and Related Measures R package version 2.6 (2023). https://CRAN.R-project.org/package=rsq.

57. Riebler, A., Sørbye, S. H., Simpson, D. & Rue, H. An intuitive Bayesian spatial model for disease mapping that accounts for scaling. Statistical methods in medical research 25, 1145– 1165 (2016).

58. Rue, H., Martino, S. & Chopin, N. Approximate Bayesian inference for latent Gaussian models by using integrated nested Laplace approximations. Journal of the Royal Statistical Society Series B: Statistical Methodology 71, 319–392 (2009).

59. Rue, H. et al. Bayesian computing with INLA: a review. Annual Review of Statistics and Its Application 4, 395–421 (2017).

60. Vaughn, R. L., Muzi, E., Richardson, J. L. & Würsig, B. Dolphin bait-balling behaviors in relation to prey ball escape behaviors. Ethology 117, 859–871 (2011).

61. Mathis, A. et al. DeepLabCut: markerless pose estimation of user-defined body parts with deep learning. Nature neuroscience 21, 1281–1289 (2018).

62. Hirt, M. R., Jetz, W., Rall, B. C. & Brose, U. A general scaling law reveals why the largest animals are not the fastest. Nature Ecology & Evolution 1, 1116–1122 (2017).

