## Supplemental Figures for "Temperature-dependent predation predicts a more reptilian future"

**Author names and affiliations:**

John M. Grady[1,2*†], Jacob L. Amme[2*], Kiran Bhaskaran-Nair[2], Varun Sinha[2], Samuel J. Brunwasser[2], Quentin D. Read[3], Sydne Record[4], Anthony I. Dell[5‡], Keith B. Hengen[2‡†]

[1]Living Earth Collaborative, Washington University in St. Louis, St, Louis, MO, USA
[2]Department of Biology, Washington University in Saint Louis, St. Louis, MO, USA
[3]United States Department of Agriculture, Agricultural Research Service, Raleigh, NC, USA
[4]Department of Wildlife, Fisheries, and Conservation Biology, University of Maine, Orono, ME, USA
[5]National Great Rivers Research and Education Center, East Alton IL, USA

[*]These authors contributed equally to this work.
[‡]Last author

### Supplemental Figures

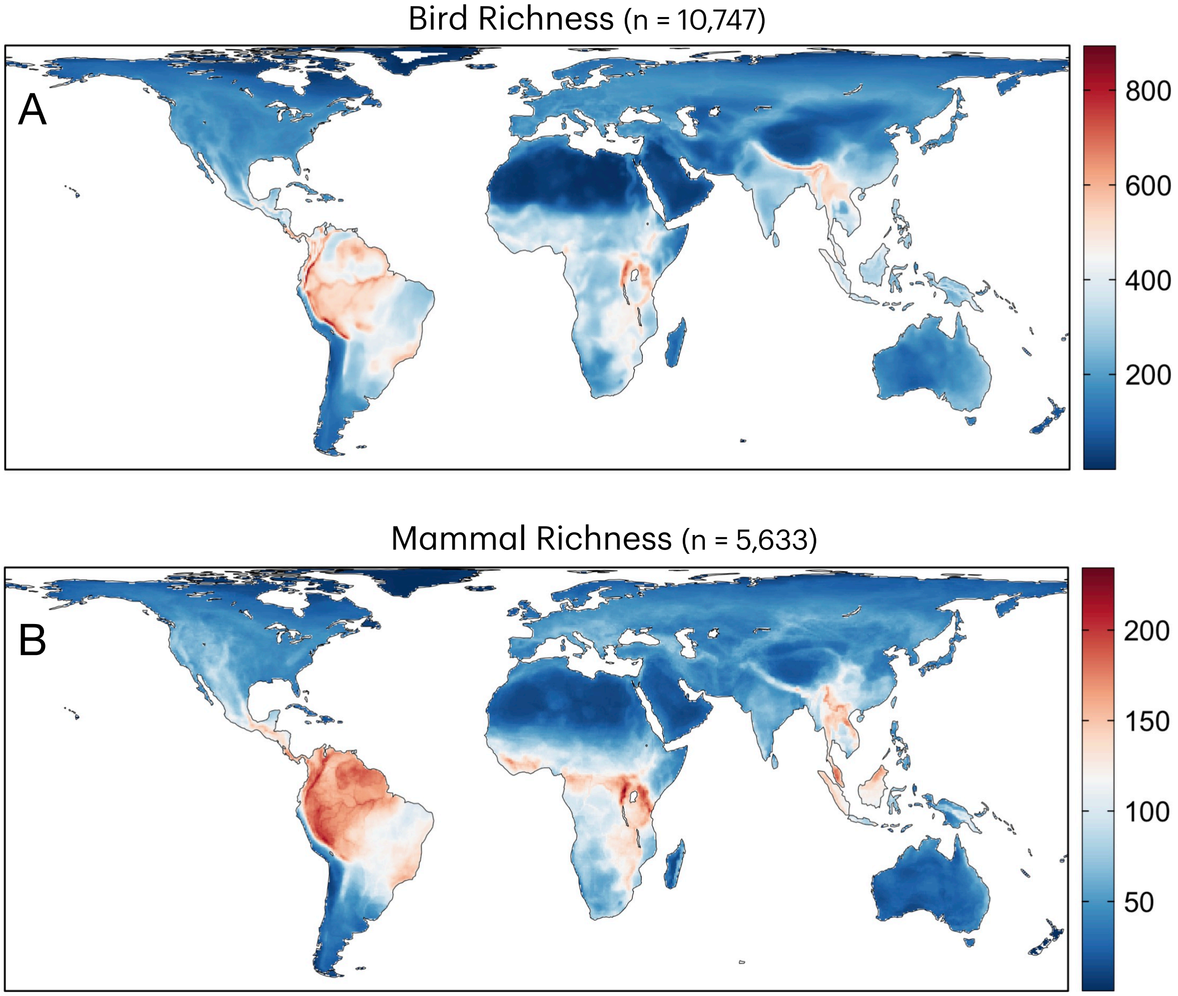


**Figure S1: Global endotherm richness for birds and mammals.** Richness was calculated by overlaying species distributional maps and summing occurrences within a 48.25 km x 48.25 km grid cell.

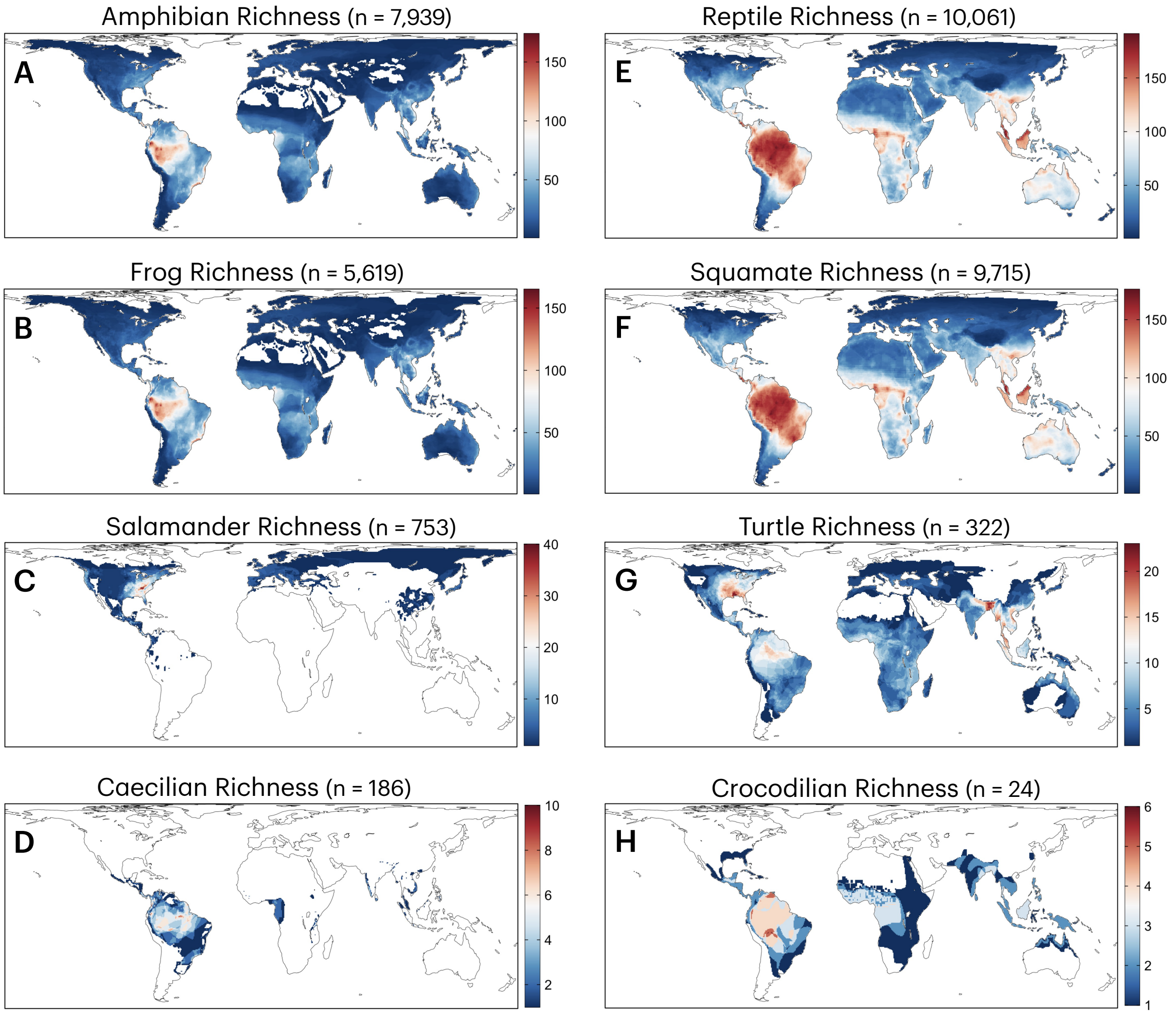


**Figure S2: Global ectotherm richness.** Richness for amphibian clades **(A-D)** and reptile clades **(E-H)**. Species n is labeled

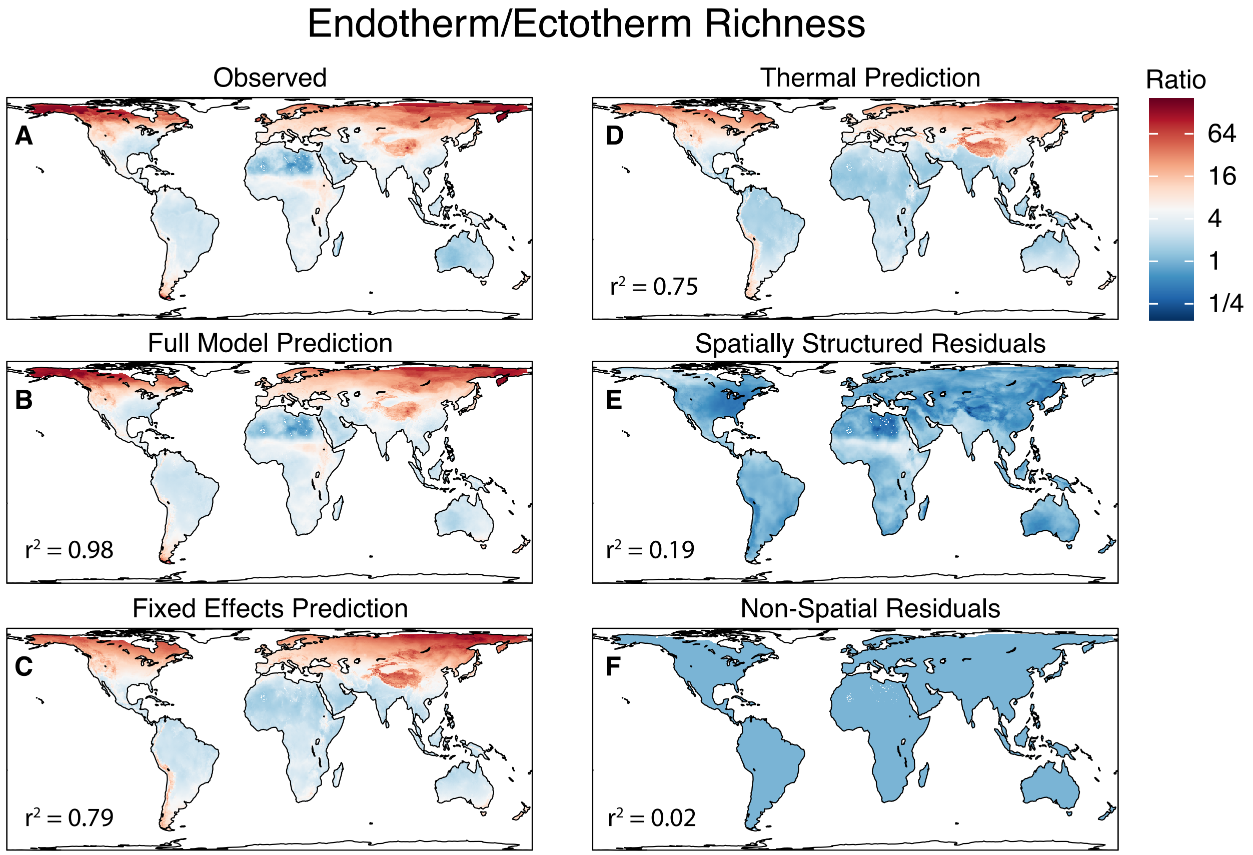


**Figure S3: Spatial model of endotherm/ectotherm richness.** **(A)** Empirical patterns of comparative richness (see also Fig. 1A). **(B)** BYM2 model predictions (Besag, York, and Mollié). BYM2 is a spatially-explicit Bayesian hierarchical model that includes main effects, spatially structured random effects, and spatially unstructured random effects. **(C)** Predicted ratio of endotherm/ectotherm richness from fixed effects alone (1/kT, NPP, precipitation, and elevation). **(D)** Predicted ratio from thermal effects alone (1/kT; T is temperature). **(E)** Spatially structured residuals (modeled spatial autocorrelation). **(F)** Non-spatially structured residuals. In E-F, the smaller range of values reflects the lower predictive power of residuals. Explained variance from variables is indicated by the $r^2$ value. n cells = 54,795

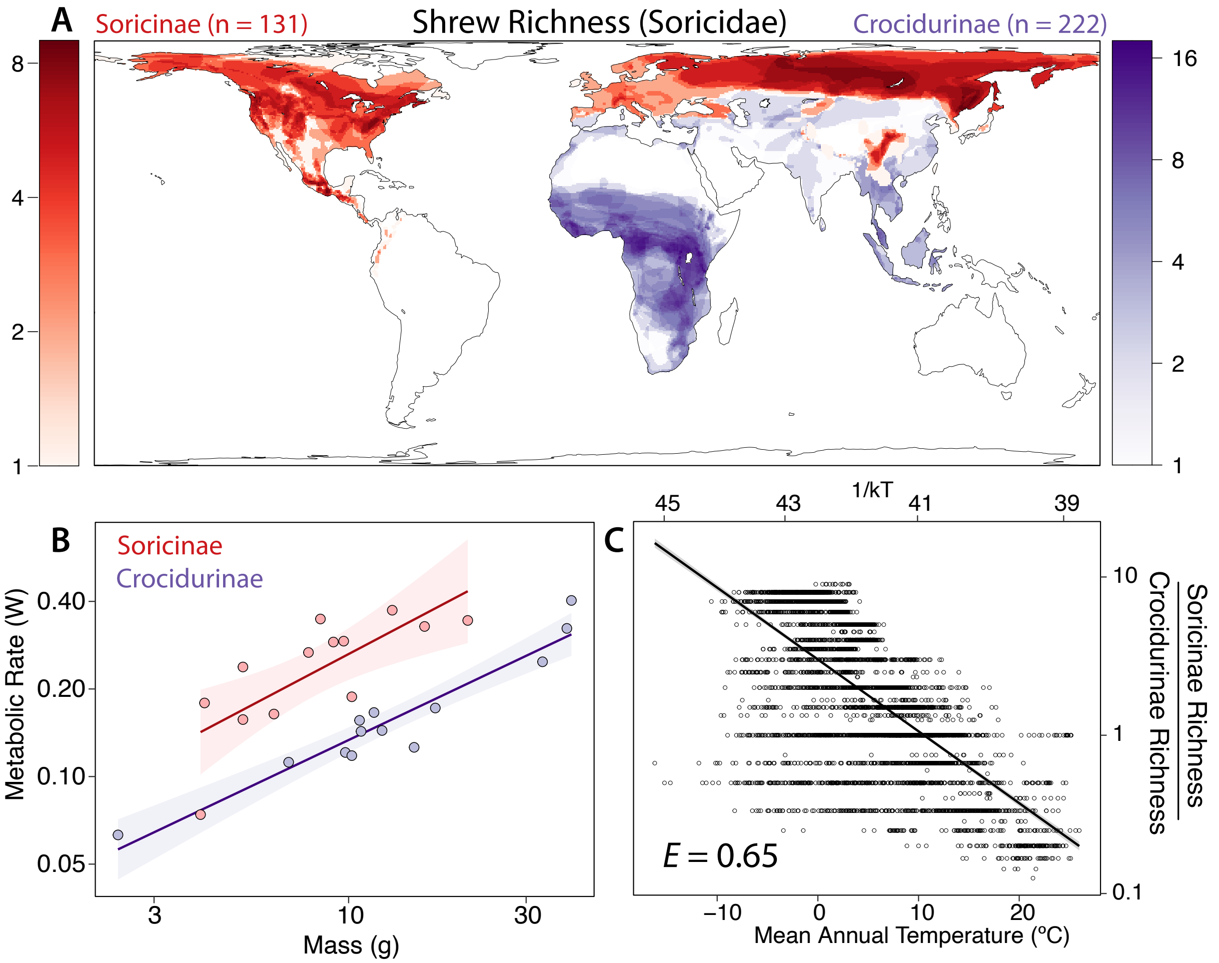


**Figure S4: Comparative shrew richness.** **(A)** Spatial patterns of two shrew subfamilies, fast-metabolizing Soricinae and slow-metabolizing Crocidurinae. These families overlap in much of central and southern Eurasia. In low latitudes, Soricinae is generally restricted to cooler, mountainous habitat, such as the Himalayas, Central American highlands and the Andes. **(B)** Basal metabolic rates by body mass in shrew subfamilies; Soricinae are approximately double Crocidurinae for a given weight. Regression: Soricinae: $y = 0.67x$–$2.89$, $r^2 = 0.57$, n = 13; Crocidurinae: $y = 0.61$–$3.41$, $r^2 = 0.91$, n =13. See Supp. Data 1 Tab 4 **(C)** The ratio of Soricinae richness to Crocidurinae richness where taxa overlap in Eurasia. Thermal sensitivity of richness: $E = 0.65$, $CI$ : 0.62–0.67, $r^2 = 0.61$ (Bayesian spatial model) is identical to the thermal sensitivity of metabolism ($E = 0.65$)[18].

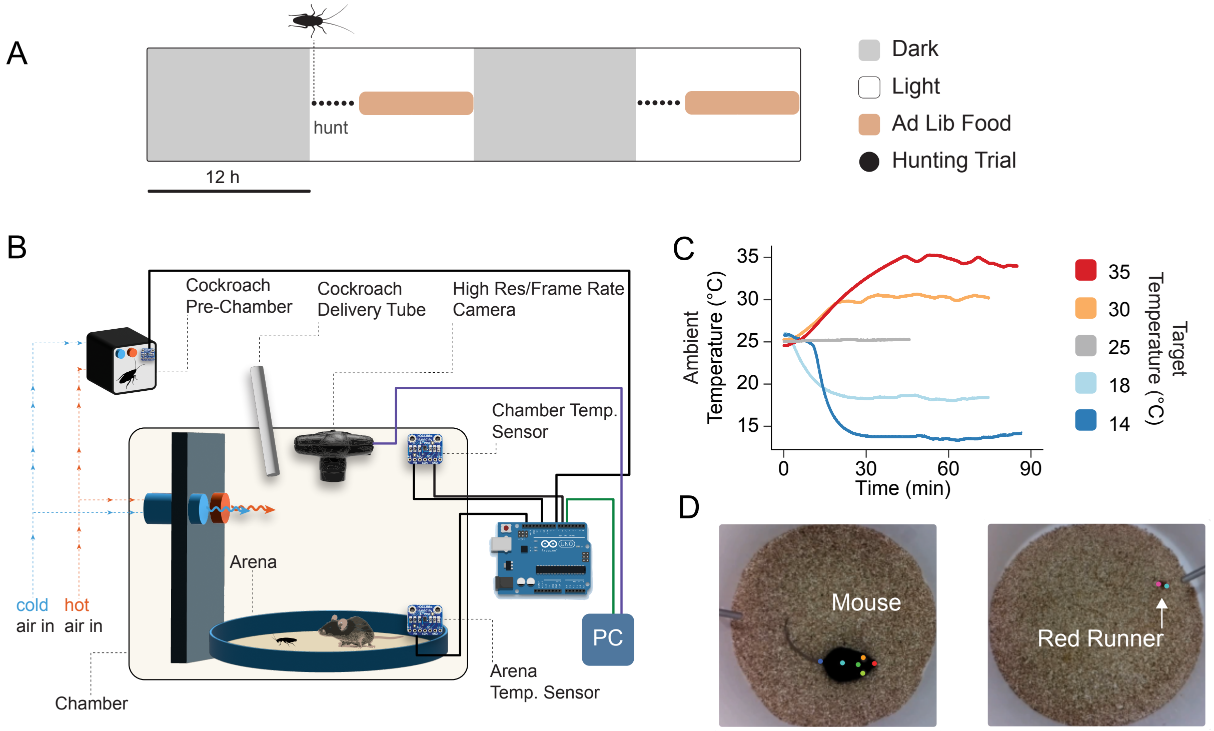


**Figure S5: Schematic of endotherm and ectotherm locomotion and predation.**(**A**) After overnight food restriction, mice were offered red runners daily, up to six trials per day. (**B-C**) Hunting arenas were heated and cooled, and animal movement was tracked (**D**) using markerless pose estimation.

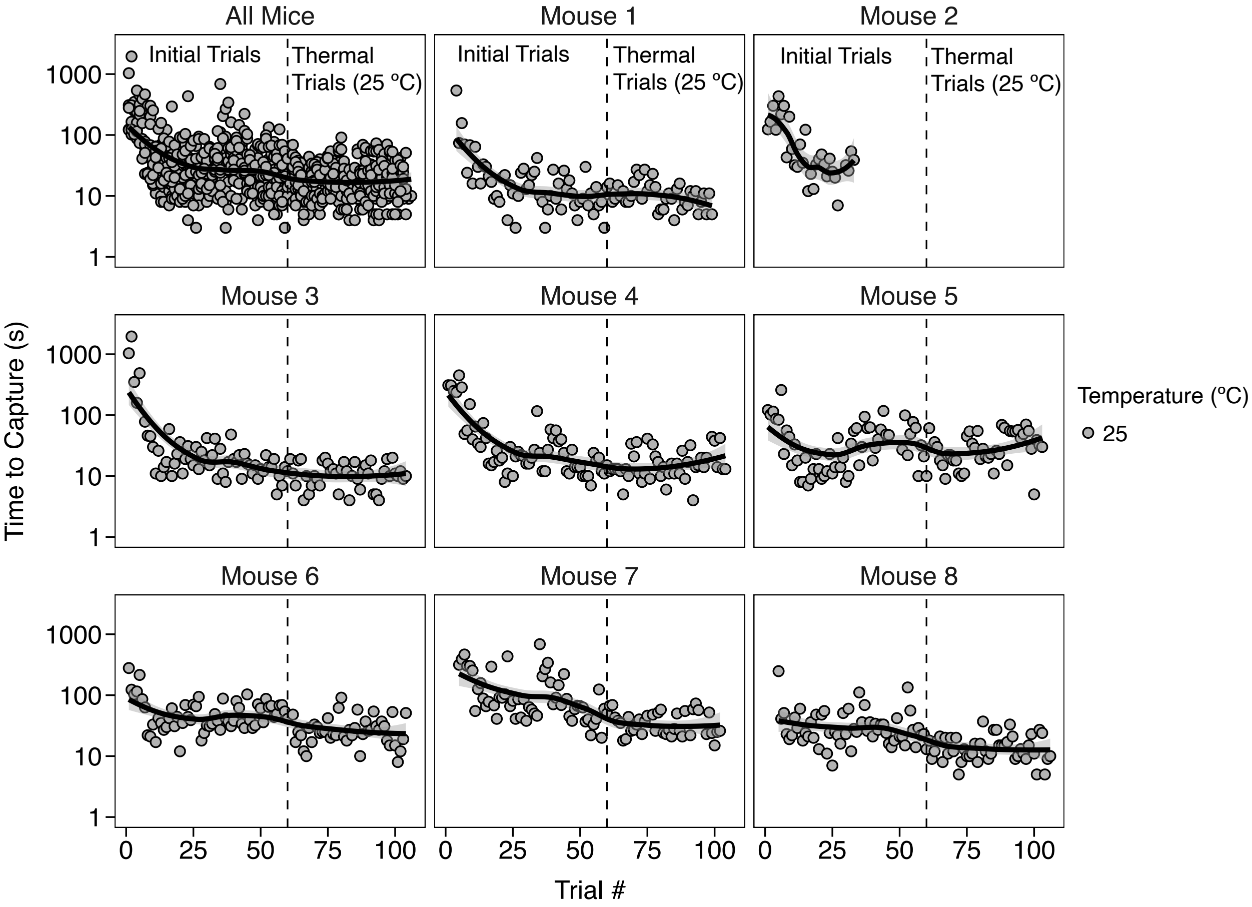


**Figure S6: Individual mouse learning curves.** The time required for mice to capture red runner prey declines with trial number, approaching an asymptote by the conclusion of initial trials (dashed line), conducted at room temperature (25 °C). Subsequent room temperature trials (right of dashed line), show only modest improvement, if any. LOESS fits are from ggplot2.

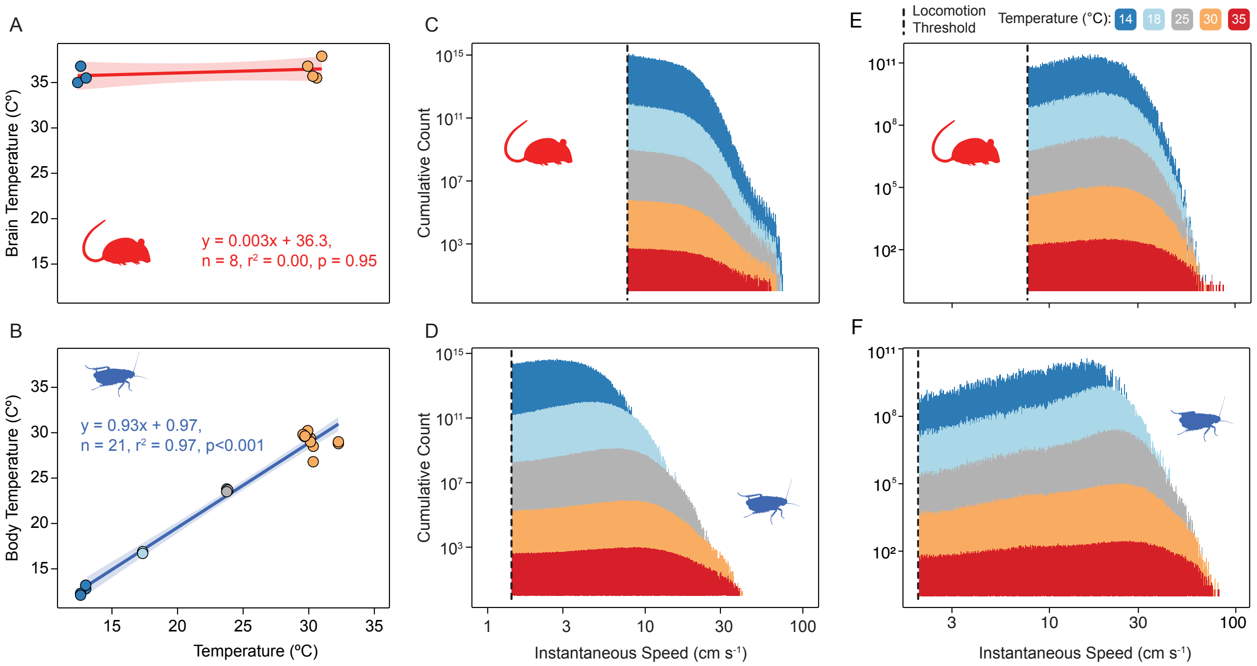


**Figure S7: Thermal sensitivity of mice and red runners.** **(A)** The internal temperature (brain temperature) of mice does not vary as a function of ambient temperature between 14 and 30 °C). Mouse brain temperature was measured by neuronal temperature probe in 8 animals (4 per temperature). **(B)** In contrast, red runner body temperature closely mirrors ambient temperature, with a slope near 1. Red runner temperature was measured by thoracic thermal probe in a total of 21 red runners across four temperatures between 14 and 30 °C. **(C-D)** Cumulative, stacked histograms of frame-to-frame speeds of mice and red runners in isolation at five temperatures between 14 and 35 °C, where each frame is 1/30 s. Y axis values refer to the sum of a video frames with movement across all thermal groups. The threshold for locomotion (vertical dashed line) represents the empirically determined velocity threshold that separates locomotion from non-locomotory movements (See Fig. S9 for thresholds). In red runners, the threshold for locomotion is lower than mice due to the absence of measurable postural adjustments at rest. Note that the distributions of mouse movement in isolation are similar at every temperature. In contrast, as temperature increases, the median and maximum speeds of red runner movement in isolation also increase. **(E-F)** Same as **(C-D)** but for mice and red runners during pursuit. Note that there is a stronger signal of temperature for red runners than for mice during a hunt.

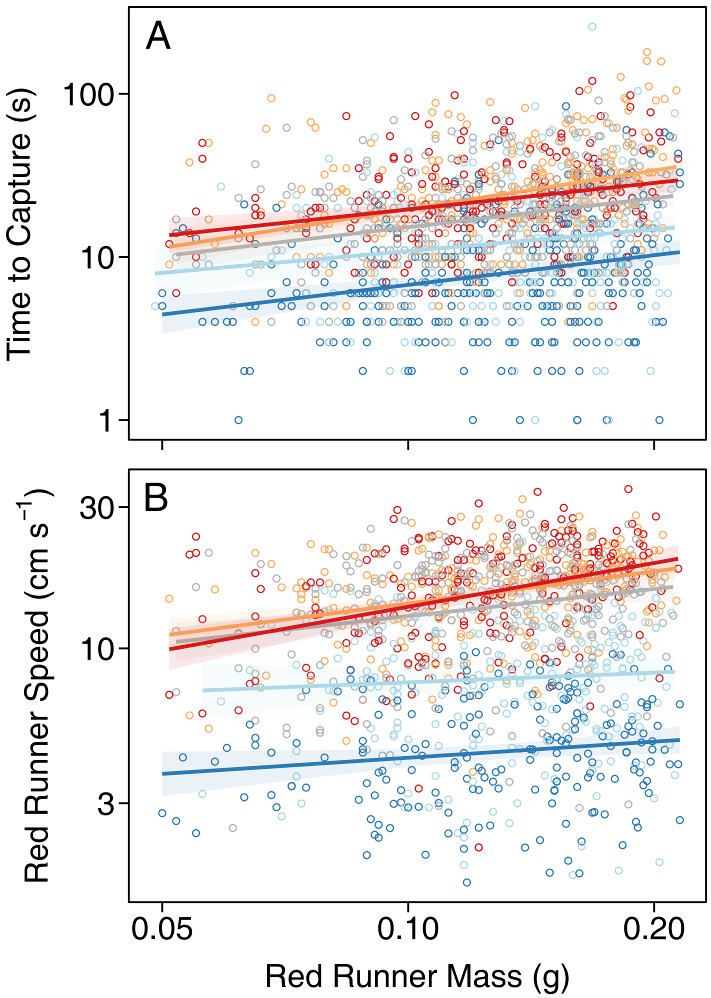


**Figure S8: The effects of red runner mass on speed and time to capture.** As red runner mass increases, the time to capture increases **(A)**, and red runner speed increases **(B)**. Color indicates thermal trial temperature, from 14 °C (blue) to 35 °C (red). To control for this effect, red runner mass was treated as a fixed effect in analyses

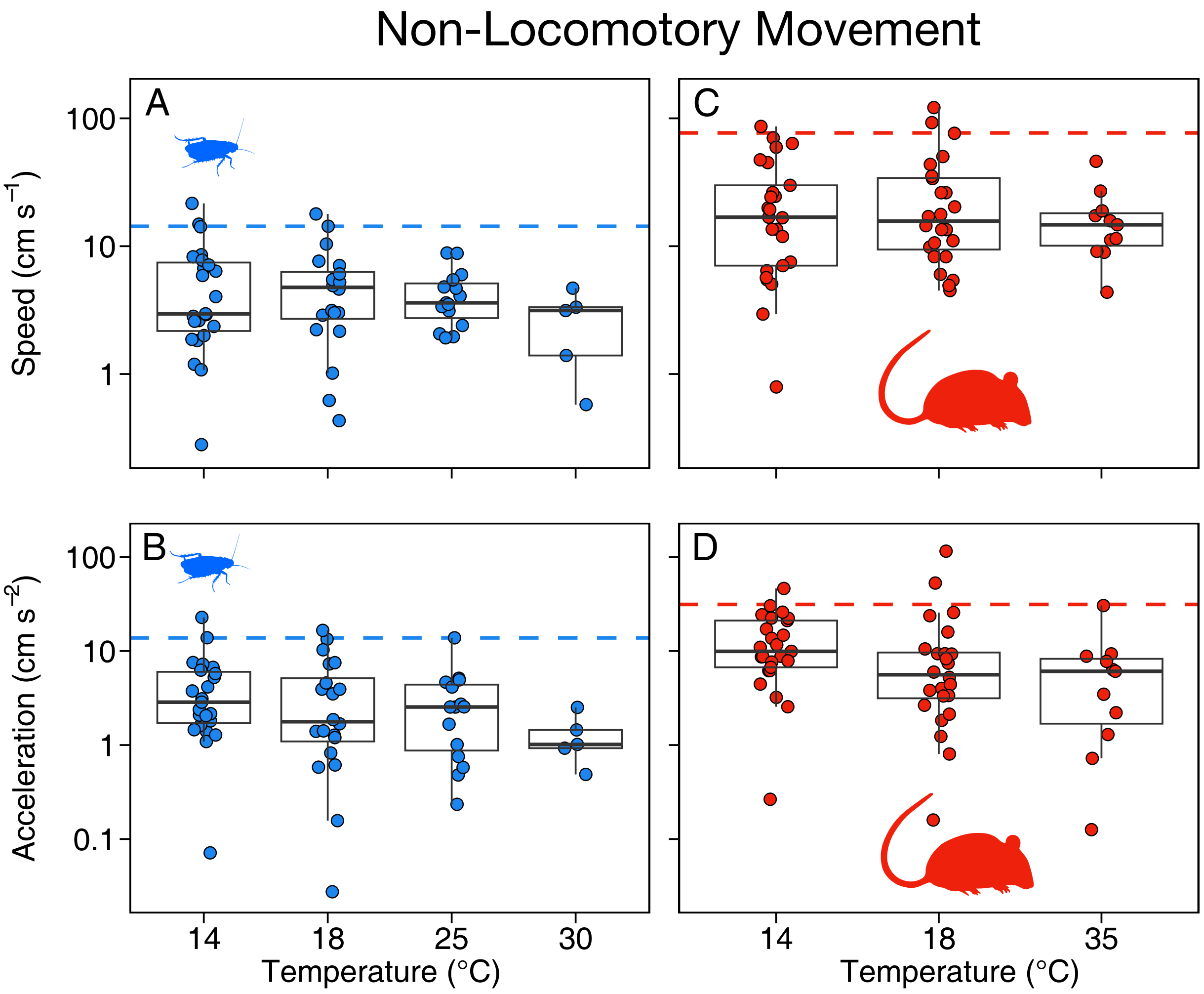


**Figure S9: Quantifying non-locomotory movement.** Videos were manually reviewed to identify frames when animals were stationary yet movement was detected. Such movement could reflect non-locomotor activities like grooming, or video artifacts caused by pixel jittering. Dashed lines indicate 95% quantiles, which were used as a threshold to distinguish locomotory from non-locomotory movement. Red runners **(A, B)** have lower thresholds than mice **(C, D)**, which exhibit more postural movements during grooming. n mouse = 60, n red runner = 63.

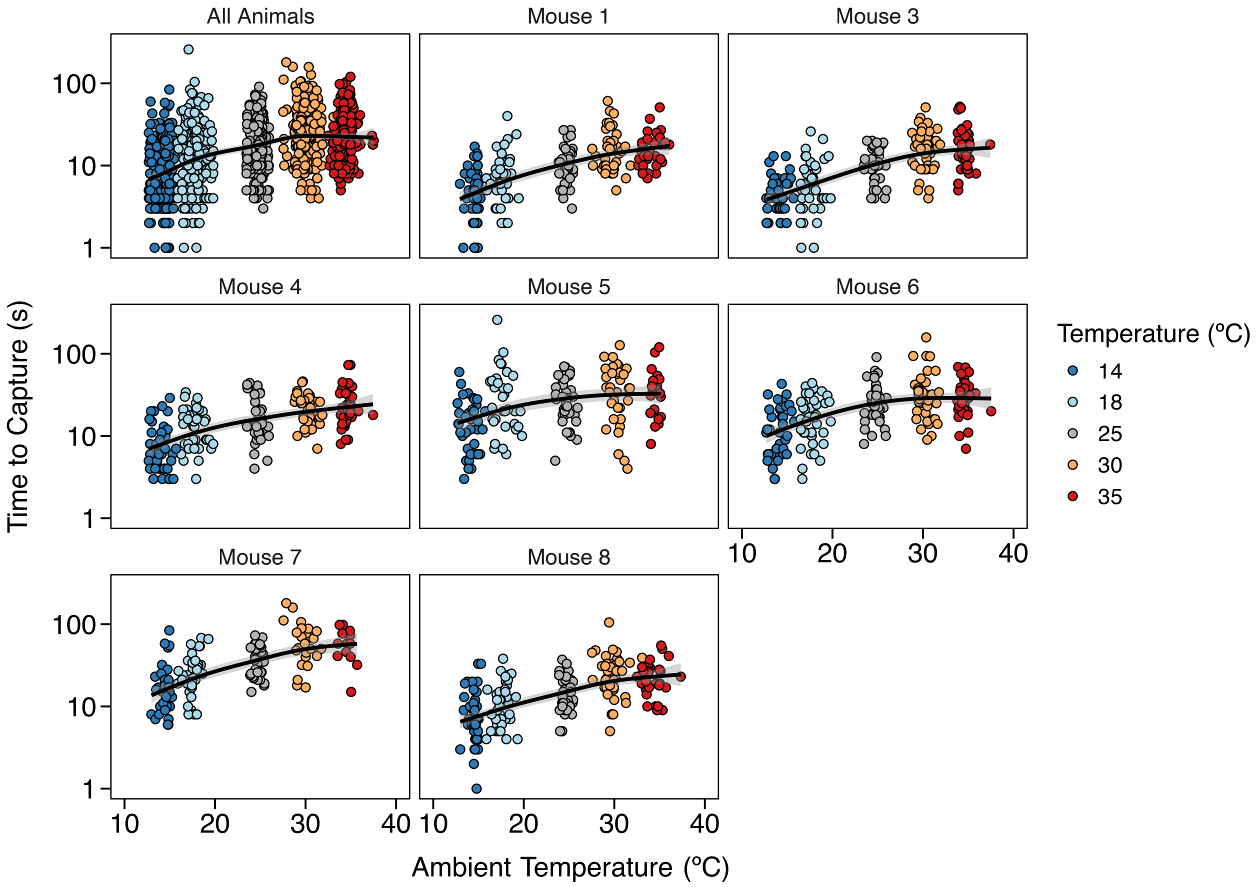


**Figure S10: Time to capture for individual mice.** When pooled, all animal show a strong thermal effect for the time required to capture prey (top left), but similar patterns are observed when considering animals separately (remaining panels). Mouse 2 stopped hunting after the initial trials, so was excluded from thermal hunting results (n mice = 7).

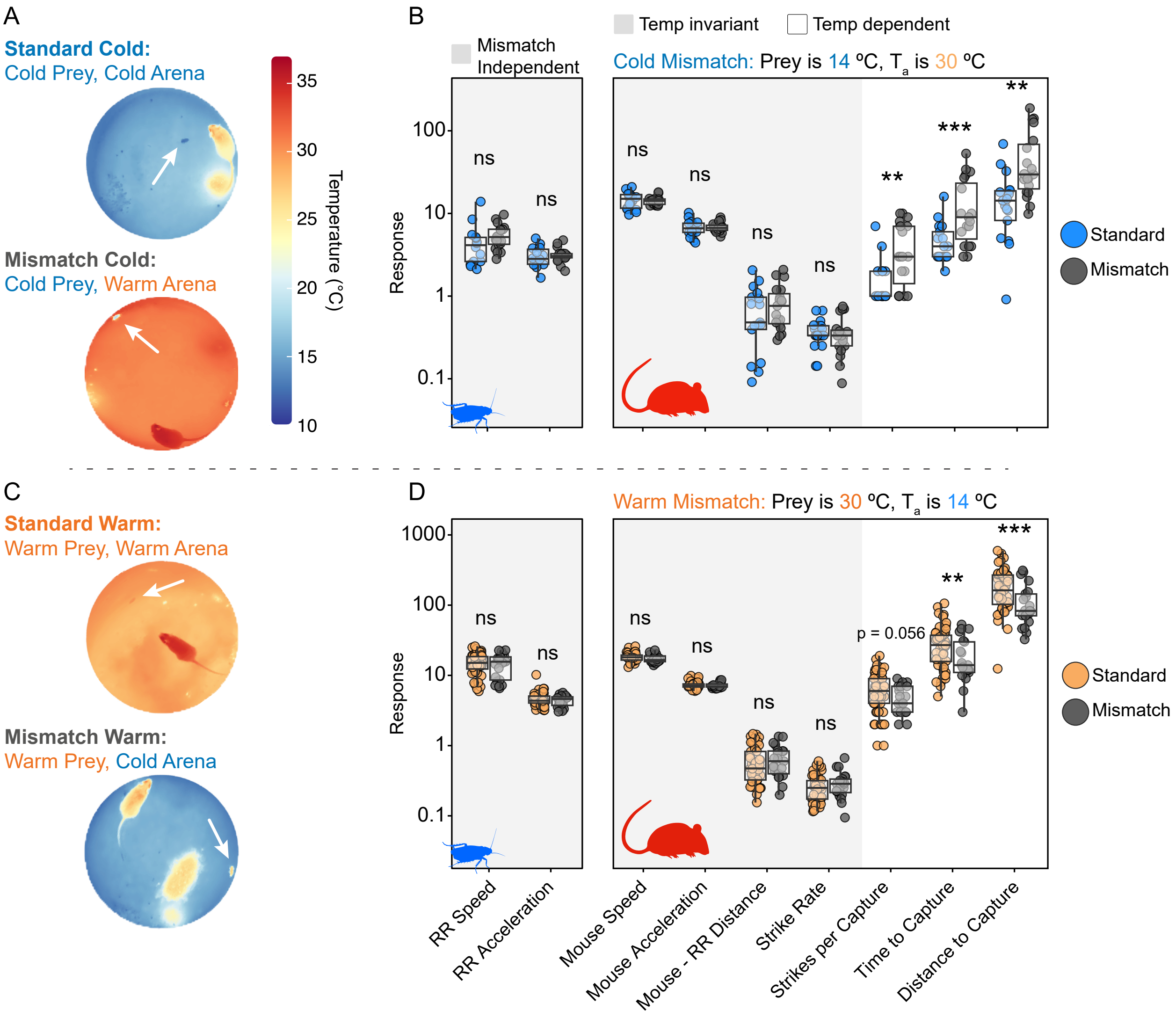


**Figure S11: Mice use ambient temperature to anticipate the behavior of future prey.** **(A)** (Top) Thermal image of hunting trials of a standard cold trial in which ambient temperature ($T_a$) and red runner body temperature ($T_b$) are both cold. (Bottom) Mismatch cold trial in which $T_a$ is warm while red runner $T_b$ is cold. White arrow indicates location of red runner. **(B)** Comparison of behavioral features as a function of condition: standard cold vs. mismatch cold. Gray background indicates features predicted to be invariant (see Fig. 5B,F). For instance, red runner body temperature is approximately the same in both conditions, so its speed and acceleration is expected be the same. If mice adjust their speed to match prey speeds, mouse locomotion should be similar between matched and mismatched conditions. Because prey distance and strike rate was previously observed to be thermally independent (see Fig. 5B,F), we also did not expect them to change between mismatch and standard conditions. Results support predictions. As expected, thermally sensitive response rates all varied between mismatched and standard conditions: strike rate, time to capture and distance traveled all showed a worse performance when mismatched (Data 1 Tab 8). **(C)** Same as (A) but for standard warm and mismatch warm trials. (Top) standard warm trial in which ambient temperature ($T_a$) and red runner body temperature ($T_b$) are both warm. (Bottom) mismatch warm trial in which $T_a$ is cold while red runner $T_b$ is warm. **(D)** Same as (B) but for standard warm and mismatch warm; however, under mismatch conditions mice performance improved (Data 1 Tab 8). Even if not perfectly adaptive, these shifts in performance with temperature are consistent with mice learning thermal cues to anticipate prey behavior. ns - not significant. $^{*}p < 0.05, ^{**}p < 0.01, ^{***}p < 0.001$. P values are for two sided Wald t test. Effect sizes associated with significant values are $\geq |0.77|$. See also Data 1, Tab 7

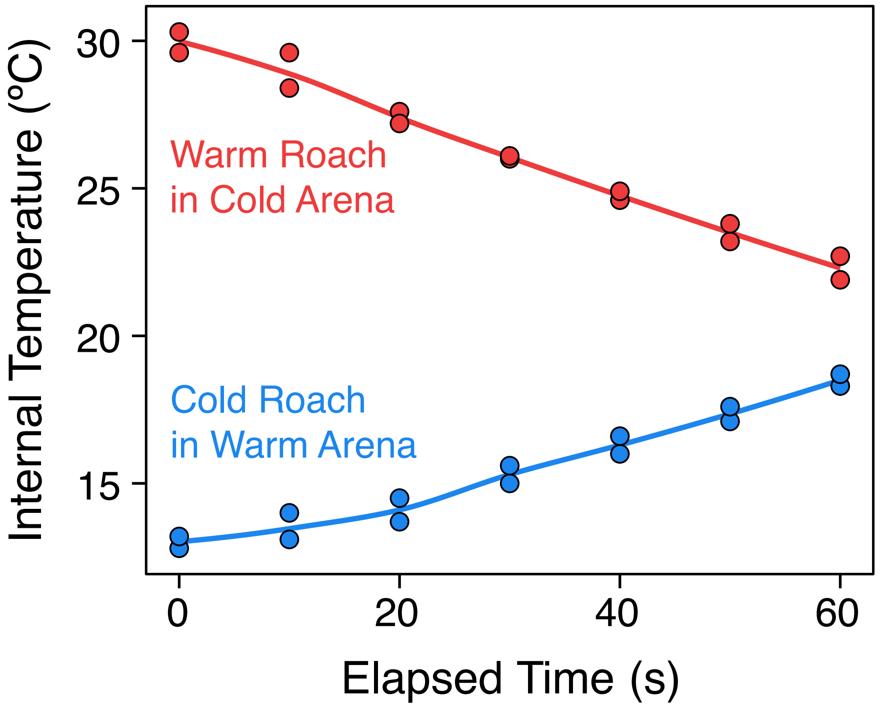


**Figure S12: Red runner internal body temperature in thermal mismatch conditions.** For mismatch trials, body temperatures initially differed from ambient temperatures; but warm roaches eventually cooled in colder arenas and vice versa. Note, however, that the time to warm or cool more than a few degrees occurs after the typical time to capture, ~3-20 s. This modest change in body temperature did not significantly affect red runner locomotory performance (see Fig. S11)

**Supplemental Data 1. Statistical results for spatial and experimental analyses. Tab 1** presents statistical outputs for Bayesian models of temperature and vertebrate diversity, which account for spatial autocorrelation. Predictor variables include ambient temperature, precipitation, net primary productivity (NPP), elevation range, as well as random spatial effects. Temperature was also considered as the only fixed effect. **Tab 2** presents similar results but for a linear model, excluding random effects. **Tab 3** presents residual normality analyses for models of species richness, evaluated separately for endotherms, ectotherms, and their richness ratio (endotherms : ectotherms). **Tab 4** shows shrew metabolic rates taken from the literature (citations included). **Tab 5** provides statistical results for experimental analysis of acceleration with temperature. **Tab 6** shows results from initial experimental trials at room temperature. **Tab 7** shows results from thermal trials. **Tab 8** shows results from thermal mismatch trials.
